# Functional Tumor Boundary and its Response to Short Mild Stimuli: First Dynamic 2D Electroimpedance Evidences of Dissipative Structures at Tissue Level

**DOI:** 10.1101/2025.01.29.635448

**Authors:** Yuri F. Babich

## Abstract

The Hanahan-Weinberg hallmarks of cancer framework conceptualizes cancer as a hostile ecosystem within the larger human system. The functional boundary of a tumor (FBT) can be characterized by the size and activity of the intermediate zone -a “battlefield”—driven by the interplay of cellular heterogeneity and environmental factors. The pioneering technology of functional imaging “Skin Electrodynamic Introscopy” has a unique combination of spatial and temporal resolution on a fairly large scanning area, which allowed dynamic visualization of the spectral impedance landscape (SEL), and thus to overlap and monitor the intermediate zone in our melanoma studies.

A new class of SEL phenomena was discovered for the first time: the test-induced manifestation of coherent dynamic macroscopic structuring at the tissue level, thus reflecting the collective processes of intercellular interactions in parameters of the intra- and intercellular level. It turned out that these effects are highly sensitive to short weak stimuli like ischemia, non-thermal EMF and MF.

Our primary objective was to identify general indicators of tumor functional boundary (FTB) manifestation within spectral electrical impedance parameters. Utilizing five datasets acquired through our innovative SEI technology, we aimed to evaluate the static and dynamic characteristics of the FTB in relation to its VB, contrasting them against the surrounding tissue.

This study interprets observed effects through the lens of the theory of self-organization and dissipative structures. The findings provide the first empirical evidence of dissipative structuring at the tissue level, observed across the majority of the investigated phenomena- as transition from initially chaotic SEL to the tumor specific patterns and back upon cessation of the stimulus:

- Initial impedance boundaries are significantly wider than visible ones, approximately repeating their shape. The stimulation leads to activation of intermediate zone and expansion of impedance boundaries, especially noticeable at the invasive fronts.
- Relaxation of intermediate zone in the form of mobile clusters;
- Emergence of antiphase structures/clusters as markers of the tumor invasive fronts,
- Emergence of the tumor resistance zone to the influence as a post effect;
- Significant spatial and temporal differences of FTB at the intra- and intercellular level, which is especially noticeable at the level of mitochondrial membranes;
- Manifestation of stroma of pre-existing nevus at the mitochondrial fluctuation field;
- Weakening or loss of the most phenomena in a case of X-rays exposure.

Additionally, we report:

- Informational significance of the ion imbalance maps;
- Common features and differences found in a papillomatous nevus;
- Antiphase structuring under stress in a plant leaf.

The results demonstrate the potential of dynamic electrical impedance imaging to delineate tumor heterogeneity and offer novel insights into cancer-host interactions during targeted interventions in real time.

**Simple Summary:** Objective assessment of tumor boundaries and its affected surrounding is a key and persistent problem for all types of cancer. Our previous in vivo experiments with melanoma firstly revealed: significant excess of impedance boundaries over visible ones and its increased sensitivity to weak and short stimuli (ischemia, electromagnetic and magnetic fields). This emerged e.g. as antiphase structuring at the visible tumor boundaries. Additional processing revealed a number of new phenomena like advance and reversal of impedance fronts in response to the stimuli. It is assumed that the revealed phenomena are the first registration of dissipative structuring at the tissue level. The results demonstrate potential of the dynamic electroimpedance imaging to delineate tumor functional boundaries and assess adaptive responses to external stimuli in real time.

## Introduction

> *If the entropy changes at the border of the tumorous and the normal tissues are compared as cancers’ velocity, then the ratio between the two forms of tissue may be characteristic of the borderline, where an overlap in entropy production rate may be found between the normal tissues and the tumor. This can be considered a new target for demarcation”. I. Prigogine*^*1*^

Although tremendous progress medical technologies has been achieved in the 21st century, early diagnosis and complete recovery of malignant tumors are still facing severe challenges: cancer remains a dominating health concern around the world. At the same time, research into cancer initiation and progression has been overwhelmingly directed towards the biochemistry, genomics, and cell biology of cancer. By contrast, far less attention has been focused on the biophysics and system biology of the cancer state^2^. Description of order and function in biological systems has been a challenge to scientists for many decades. The overwhelming majority of biological order is functional order, often representing self-organized dynamical states in living matter^3^. Systems biology has emerged as a powerful paradigm in the life sciences. It is based on the assumption that the properties of complex systems consisting of many components interacting with each other in a nonlinear, nonadditive manner cannot be understood by focusing only on the components. The system as a whole has emergent properties that *are not visible at the level of its parts but o*nly at the *macro view*^4^. The process of self-organisation is often triggered by *seemingly random fluctuations*, amplified by positive feedback. Examples include crystallization, cyclones, Zhabotinsky reactions, all living matter. Ecological examples presupposes spatial and functional differentiation like bacteria and ant colonies, flocking behaviour etc.

The physics of this phenomenon is based on the theory of dissipative structures (DS), a concept introduced by Ilya Prigogine, a Nobel laureate in chemistry, to describe systems that maintain themselves in a state far from equilibrium by dissipating energy. DS are characterized by their ability to exchange matter, energy, and information with their environment to sustain their existence, they occur when a steady state becomes unstable at a critical bifurcation point^6^. DS exist at all scales, systems, and at different levels of complexity: from the universe to quantum mechanics^7,8^. Recent research into bio-analogue dissipative electrically and chemically driven systems revealed that some of these structures exhibit organism-like behavior, reinforcing the earlier expectation that the study of DS will provide insights into the nature of organisms and their origin^9-11^.

The Hanahan-Weinberg hallmarks demonstrate cancer is a hostile ecosystem within a larger open human system: a neoplasm is formed on the site and from parts of a previous system, which found itself at a bifurcation point^5^. The intermediate zone between these systems represents the battlefield where the tumor and the host engage in a complex interplay of interactions. The tumor attempts to exploit the host’s resources and evade its defenses, while the host attempts to eliminate the tumor. This interaction involves a constant exchange of signals, including growth factors, cytokines, and metabolic products, which shape the tumor’s behavior and the host’s response. The outcome of this competition determines the tumor’s fate and the overall health of the host.

The predominant focus in cancer research has been on the tumor microenvironment (TME) operating at the micrometer scale, largely due to the accessibility of cellular and molecular components for study. In contrast, the dynamic functional boundaries of the tumor (FBT), extending well beyond the micrometer-scale TME, have received considerably less attention. This discrepancy is primarily due to the significant challenge of visualizing the tumor functional landscape in real time^84^. Understanding these broader boundaries involves understanding how the tumor interacts with and influences distant tissues and systemic physiology, which are difficult to capture with current imaging techniques, which often provide static snapshots rather than dynamic views^85^. The lack of tools to monitor these larger-scale dynamic interactions in real time limits the ability to fully characterize the functional consequences of expanding tumor influence and its systemic effects^86^.

The assessment of FBT is a critical component in the diagnosis, treatment planning, and monitoring of cancer progression. Traditional methods for delineating tumor margins often rely on static imaging techniques such as MRI or CT scans, which may not provide an accurate representation of the tumor’s dynamic behavior or its interaction with the surrounding tissues. This limitation underscores the need for innovative approaches that can visualize and monitor tumor boundaries in real time, both in vivo and in vitro. By identifying regions where healthy tissue interacts with tumor cells, researchers can explore novel treatment strategies aimed at disrupting these interactions or restoring normal tissue function^12^.

There are few references (and, apparently, nothing on bioelectroimpedance) in the literature to in vivo tissue studies on visualization of DS effects at tissue level. To distinguish malignancy from benignity, current clinical and laboratory methods primarily rely on classical thermodynamics of equilibrium processes (volume, temperature, pressure) rather than nonequilibrium ones^13^. Common methods for tracing dissipative self-assembly, such as dynamic light scattering, electron microscopy, and rheological methods, have some drawbacks, including signal delay, low sensitivity, and low spatial and temporal resolution. The fluorescence method is more popular due to its low cost, ease of use, high sensitivity, and real-time monitoring capabilities^14^.

However, it also has some known limitations: dependence on probes, quenching, and potential damage from both light exposure and the fluorophores themselves. Given the dynamic nature of dissipative structures, their direct visualization and real-time monitoring are highly desirable as they can provide ‘movies’ rather than merely ‘snapshots,’ offering a better understanding of the dynamic aspects^15^.

Intercellular interaction is key to the homeostatic life of multicellular communities. From the moment a cell is on the path to malignant transformation, its interaction with other cells from the microenvironment becomes altered^87^. All physiological phenomena, specifically communicational, directly or indirectly, are inherently bioelectrical^16^. Therefore, electrical impedance spectroscopy (EIS) appears to be particularly promising and, in recent decades, became already established technique in both in vivo and in vitro cancer studies. In general, the advantages of electrical methods for studying information processes, including intercellular interactions, are their ability to provide real-time data with high resolution, sensitivity to subtle changes, and non-invasiveness when applied to intact biological systems^17^.

Electrometric methods in general and electrical impedance ones in particular, has become a powerful tool, allowing for rapid, non-invasive, and label-free acquisition of electrical parameters of single cells^18^. These studies have long been a mainstay of cancer research, providing controlled environments to observe cell behavior. However, these models often fail to replicate the complexity of living tissue. The limitations inherent in two-dimensional cell cultures or even three-dimensional spheroids highlight the necessity for more representative models and technique that can capture the dynamic interplay between cancer cells and their microenvironment. However, the existing in vivo EIS has largely remained a static method that does not provide real-time visualizations. To study heterogeneous dynamic structures in biological tissue, it is essential to incorporate adequate spatial and temporal resolution into time-lapse impedance assays. Conventional EIS have only partially bridged the gap between the macro- and microscale measurements: its temporal resolution remains in the range of several minutes, rendering it unsuitable for imaging metabolic landscape with rapidly changing properties^19-23^. Thus, the delineation of real tumor boundaries is a common problem for all methods, like optical or conventional EIS, based on assessment of the tissue stationary features. Due to the lack of objective criteria, surgeons follow unfounded instructions to make indentations from the visible borders of melanoma (VB), risking both removing healthy tissue and leaving malignant one.

The concept of negentropy/information flow between cancer and healthy cells is an intriguing topic of research in cancer biology. Recent discoveries suggest that tumors may exhibit self-organizing properties influenced by their surroundings. This concept posits that as tumor cells proliferate, they create gradients of biochemical signals and mechanical forces that shape their local environment. These gradients can lead to distinct functional boundaries around tumors—areas where cellular behavior transitions from normal to malignant^24,25^.

It has been established that the rate of entropy production of cancer cells is always higher than that of healthy cells, but there is one interesting exception: provided that no external field is applied to them. When an external force field like MF and EMF is applied, the rate of entropy production of normal cells can exceed the rate of entropy production of cancer cells. Thus, it was concluded that reversing the flow of entropy in coexisting normal and tumor tissues can arrest tumor development due to reverse signal transduction in the tumor-host entity^26,27^. Despite these promising theoretical insights, practical implementation remains hindered by significant challenges. One major limitation is the current lack of adequate imaging modalities capable of visualizing DS in real-time. Such imaging techniques are essential for providing biofeedback during experimental interventions, allowing researchers to monitor changes at cellular and systemic levels, and adjust treatment protocols accordingly.

The interplay between an electrically charged tumor and its surrounding cellular environment can be understood through the lens of electromagnetism and dissipative structures. As a tumor exhibits unique electrical properties^28-31^, it alters the state of nearby conductive mediums, including adjacent cells and intercellular fluids. This alteration manifests as a dynamic SEL that reflects the tumor’s metabolic activity and heterogeneity. When external stimuli interact with this environment, they can induce modifications in the tumor’s metabolism, further influencing its electric field. Consequently, the changes in electrical parameters not only signify the tumor’s characteristics but also reshape the SEL around it, creating a feedback loop where weak external effects can lead to significant alterations in both tumor behavior and its surrounding cellular milieu.

Imaging of dissipative structuring in cancer research is crucial for understanding the complex dynamics of tumor behavior and progression. This imaging can provide valuable insights into how cancer cells interact with their environment, adapt to therapeutic interventions, and develop resistance. Effective visualization is essential for translating these insights into targeted therapies, as it enables researchers to identify critical patterns and mechanisms that drive tumor evolution and response to treatment. The fact that tumors grow over months or years while cellular responses can occur within minutes underscores the need for a real-time approach to studying these processes. By employing techniques such as skin electrodynamic introscopy (SEI), researchers can gain insights into how tumors communicate with their environment, particularly during critical moments when they respond to stimuli such as hypoxia or EMF.

Skin, the largest organ of the human body, serves as a critical interface between the internal physiological processes and external environmental factors. Its complexity and dynamic nature make it an intriguing subject not only for dermatologists but also for oncologists and physicists. The study of skin, particularly in relation to melanoma offers insights that extend beyond traditional molecular biology. Unlike other aggressive tumors, melanoma has a visually distinguishable morphological border. This distinct feature provides unique opportunities to study the spatio-temporal dynamics of this ecosystem within a living organism. By examining this macro-region through the lens of DS theory, researchers can gain a deeper understanding of tumor behavior and functional boundaries of this battle.

The advent of SEI marked a significant leap in dynamic/functional skin imaging. For the first time globally, back in the late 80s, SEI provided adequate visualization of the skin electrical impedance landscape (SEL), first in statics and then in dynamics. The innovation was particularly crucial in identifying electrophysiological portrait of melanoma at the background of visibly normal tissue^32-35^. In a series of experiments, it was established that in response to weak and ultra-weak stimuli, on and beyond the boundaries of melanoma, i.e. on visually normal skin, the surrounding landscape changes dramatically by emergence of high-amplitude coherent dynamics as in-phase and anti-phase structures, and wave-like processes.

SEL of healthy skin is distinguished by the relative smooth chaotic landscape with low-amplitude fluctuations (Fig. 1a), and its high resistance/imperturbability to the used sub-threshold test stimuli mechanical, electrical, EMF, MF, hypoxia etc., which allows it to be defined as an equilibrium system. The term “chaotic nature” in the context of skin refers to the complex and dynamic interactions that occur within the skin at various levels, including cellular and molecular processes. “When a system is far from equilibrium, small islands of coherence in a sea of chaos can appear. These structures are fragile; they last for a while, and then disappear, making way for new ones^36^”. In the case of a tumor such as melanoma, disruptions in the normal regulatory mechanisms of the skin can lead to a loss of coordination among different cell types, altered signaling pathways, increased sensitivity to external and internal factors. As all our experiments with melanoma (11) have shown, the IB significantly exceed the optical ones. Moreover, they can also differ significantly in intra- and intercellular parameters. This suggests that a comprehensive assessment of tumor boundaries must consider varying scales of tissue examination to accurately delineate the peritumoral area. The FBT are understood to be not stationary, but probabilistic dynamic limits within which the tissue demonstrates altered intercellular communications in the form of an increased sensitivity/response to mild and short stimuli such as ischemia and EMF, thus indicating its weak tumor-modified dysfunction.

**Fig. 1.**
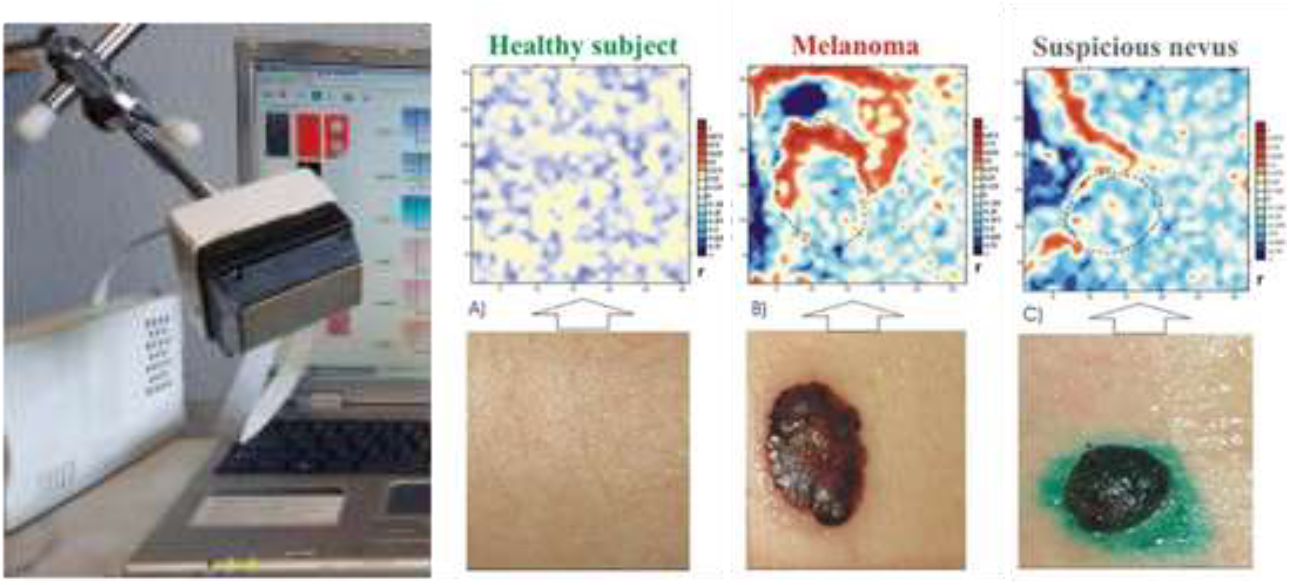
Photo of the experimental setup of SEI and examples of SEL in normal and neoplastic areas demonstrating emergence of antiphase structures around tumor invasive fronts in response to nonthermal mm-EMF [35].

By mapping out the SEL response <u>to short and weak stimuli</u> across different regions of the skin dynamically, it becomes possible to visualize areas where significant deviations occur—indicative of the tumor inhomogeneity and functional spectral boundaries of real and potential malignancy.

Given the above theoretical paradigm and our experimental findings, we can substantiate the following hypotheses:

- Among the various heterogeneous tumor segments exhibiting fluctuating states between disorder and order in response to external stimuli, there are those that are most metabolically active (e.g. younger tissue). These invasive fronts can be more briefly designated as emitters of entropy or negentropy*;
- The initial electroimpedance portraits of melanoma reflect quasy-stationary approximate flow boundaries of these emitters flow within tissues influenced by both prolonged oncogenic processes (spanning months to years) and immediate metabolic activities;
- The minimum FBT should be assessed based on its maximum IB. The width of this dynamic zone can be determined in response to weak energy or information impact: the contour line of sharp weakening of SEL oscillations is a demarcation indicator between healthy and tumor-affected tissue;
- The shape of the field/flow from a large (non-point) tumor is defined as a superposition of its emitters;
- Mitochondria** can play a primary role in establishing DS that shape the SEL and thus define its own - spectral - dimensions of FBT.

*\*The idea of defining tumor parts or boundaries as emitters of entropy or negentropy hinges on understanding how these regions respond to external impacts. If the impact leads to increased disorder within the tumor cells—then one could argue that these regions are emitting entropy. Conversely, if certain parts of the tumor respond adaptively by organizing cellular processes or enhancing survival mechanisms under stress, they might be viewed as emitters of negentropy*

*\*\* Mitochondria are often referred to as the powerhouses of the cell because they generate ATP through oxidative phosphorylation, a process that directly relates to negentropy*.

Data from 4 early experiments (3 melanomas and papillomatous nevus) were used for detailed analysis due to their similarity in the striking predominance of the IB over the visible ones. To further illustrate the dissipative nature of the discovered phenomena, we also briefly present results from a recent experiment conducted on a plant tissue.

## Methods

The methodology used was described in our previous publications^30,31^ Here we present some additional considerations in connection with the above hypothesis about the discovered phenomena as DS manifestations.

Briefly, SEI in contrast to conventional/stationary EIS, is a novel dynamic imaging modality. This is ensured by such technical advantages as unprecedented combination of three mutually contradictory features: high spatial and temporal resolution (1mm, 4ms/pixel totally for all 4 spectral parameters) over a fairly large area). That is, in time-lapse imaging mode:

- the time resolution is 4 sec and 8 sec for a frame of 32×32 mm^2^ and 32×64 mm^2^, respectively;
- spectral impedance parameters Z=|Z|e^jφ^: |Z_k_|,φ_k_, |Z_M_|, φ_M_, where j is the imaginary unit, |Z_k_|,|Z_M_| (further without parentheses), and φ_k_, φ_M_ represent the modulus and phase at 2 frequences: 2kHz and 1MHz, respectively. These parameters reflect, simplifying, the resistive (ionic) and capacitive (membrane resistance/integrity) parameters of the extra- and intracellular fluids (ECF, ICF), respectively.

The following were used as short-term stimuli: ischemia, extremely low-energy mm-EMF with low-frequency modulation and weak MF with a change in polarity (in the interval between frames to eliminate interference) as an imitation of ultra-low-frequency MF. In EIS studies of melanoma, the ratios of the spectral impedance parameters are also used as local (point) diagnostic criteria^39^, for example, k= Z_k_/Z_M_ or Z_k,,n_-Z_M,n_ (where “n” means normalised values). In the context of spatial imaging and the lack of a generally accepted term, we would define it as the “Ion imbalance index” (IB): a 2D dimensionless morpho-functional parameter, the understanding of which seems especially important in its 2/3D use^34^.

To assess FBT, both correlation and fluctuation field methods were used, since they provide complementary information. Correlations can identify regions where cellular activities are in sync or out of sync—indicating potential areas of collective behavior or vulnerability—while fluctuations reveal heterogeneity that could signify adaptive resistance mechanisms or localized variations in treatment response.

By combining both approaches, one can gain a more complete understanding of tumor dynamics under energetic or informational stress conditions such as ischemia or mm-EMF, ultimately providing insight into reprogramming strategies that target both coherent structures and fluctuating behavior within tumors.

Importantly, when analyzing fluctuation or correlation fields, we record averaged dynamics that reflect coherent fluctuations in ensembles of thousands of cells. Accordingly, when discussing various intra- and intercellular processes, we refer to their average or collective characteristics, which represent the state of the intercellular communication n network, as opposed to the generally accepted terms used in single-cell biology.

## Results and Discussion

### *<u>Analysis I</u>*. Melanoma with satellite^34^ (MS) under ischemia stress

Here we present the results of a more thorough analysis of experimental data on previously discovered but not fully understood phenomena of the SEL test-induced structuring. The previously described experiment^34^ included monitoring the melanoma zone in response to a short-term ischemia by clamping the femoral artery with a cuff: before (2.5 min), during (1.5 min) and in the process of reoxygenation (4.5 min). Time resolution - 8 sec.

Fig. 2 schematically shows the IB and zones of the MS, the statics and dynamics of which are shown in Figs. 3-5. There, unlike the visible image, not only external but also internal structures emerge, e.g.:

**Fig. 2.**
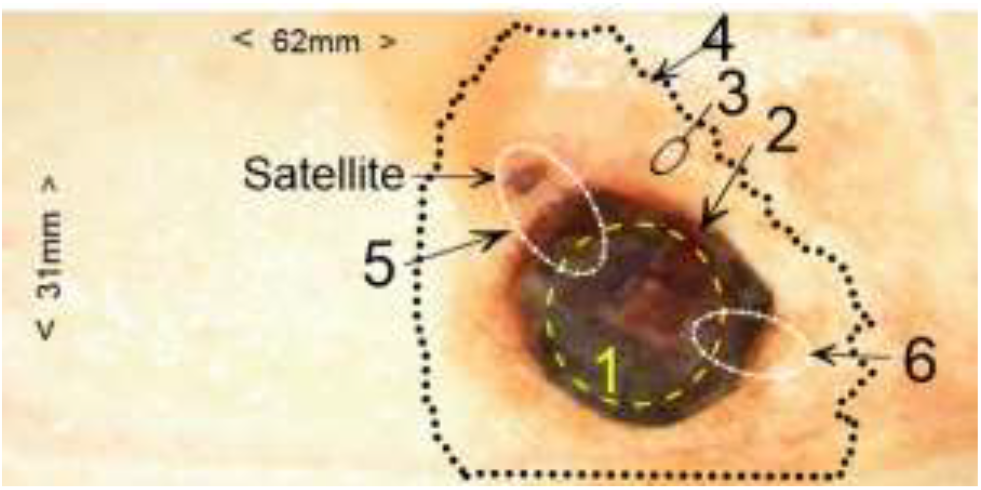
Schematic representation of the MS borders and zones. 1 -contour of the pre-existing nevus; 2- visual boundary; 3 –intermediate zone; 4- impedance boundary (by φ_k_); 5- invasive front zone with satellite; 6- invasive front zone with a satellite that has presumably not yet emerged.

**Fig. 3.**
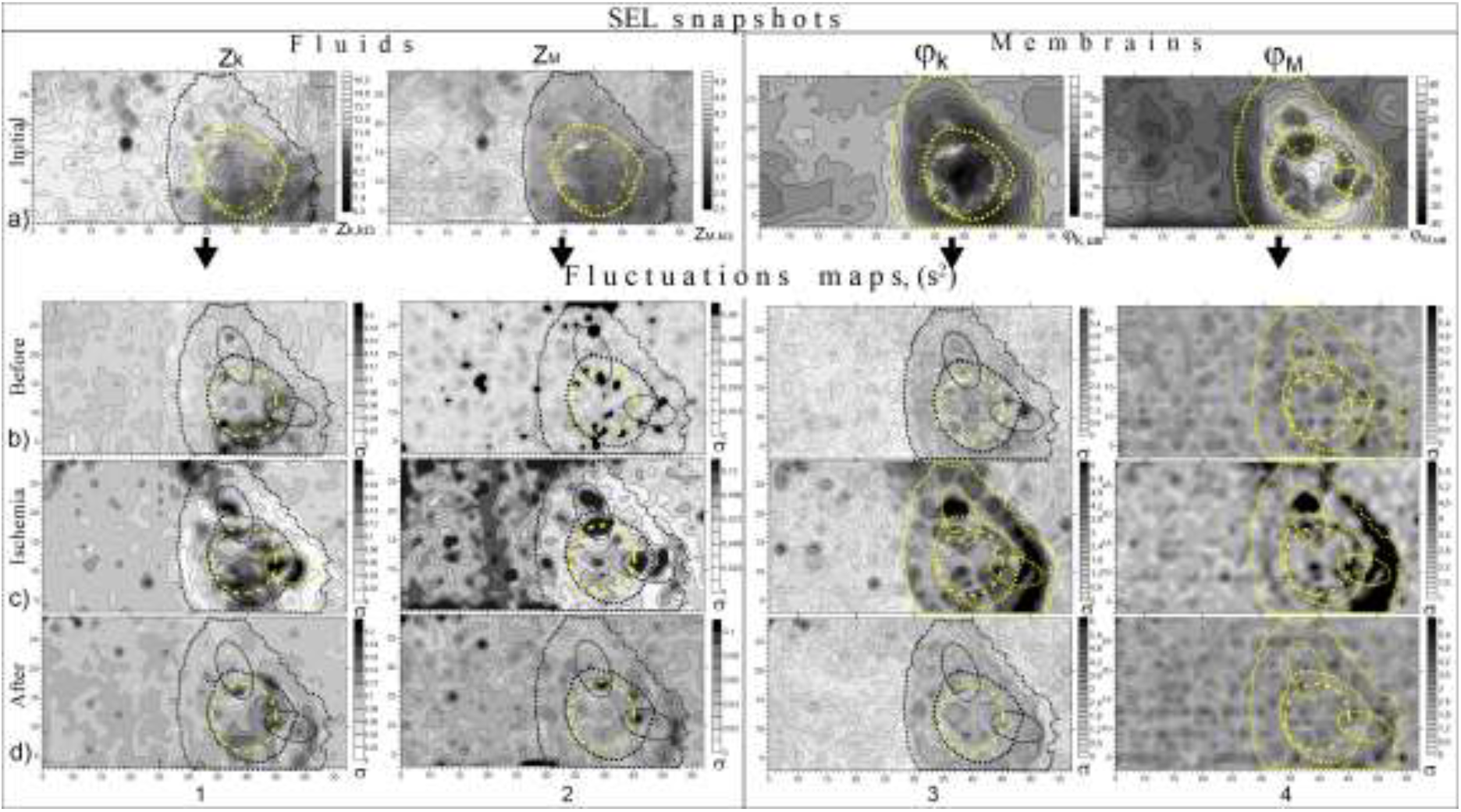
From chaos to order and back. **a)** The initial SELs in four extra- and intracellular parameters: Z_k_, Z_M_, φ_k_, φ_M._ **b-d)** The fluctuation maps calculated for the initial (b), ischemic (c) and reoxygenation (d) periods (of 15 frames, i.e. 2.5 min each). ***Note***: *The satellite and PN zone are displayed in contrast in the snapshots only on the mitochondrial landscape φ*_*M*_, *the topology and dynamics of which are the focus of the analysis below*

**Fig. 4.**
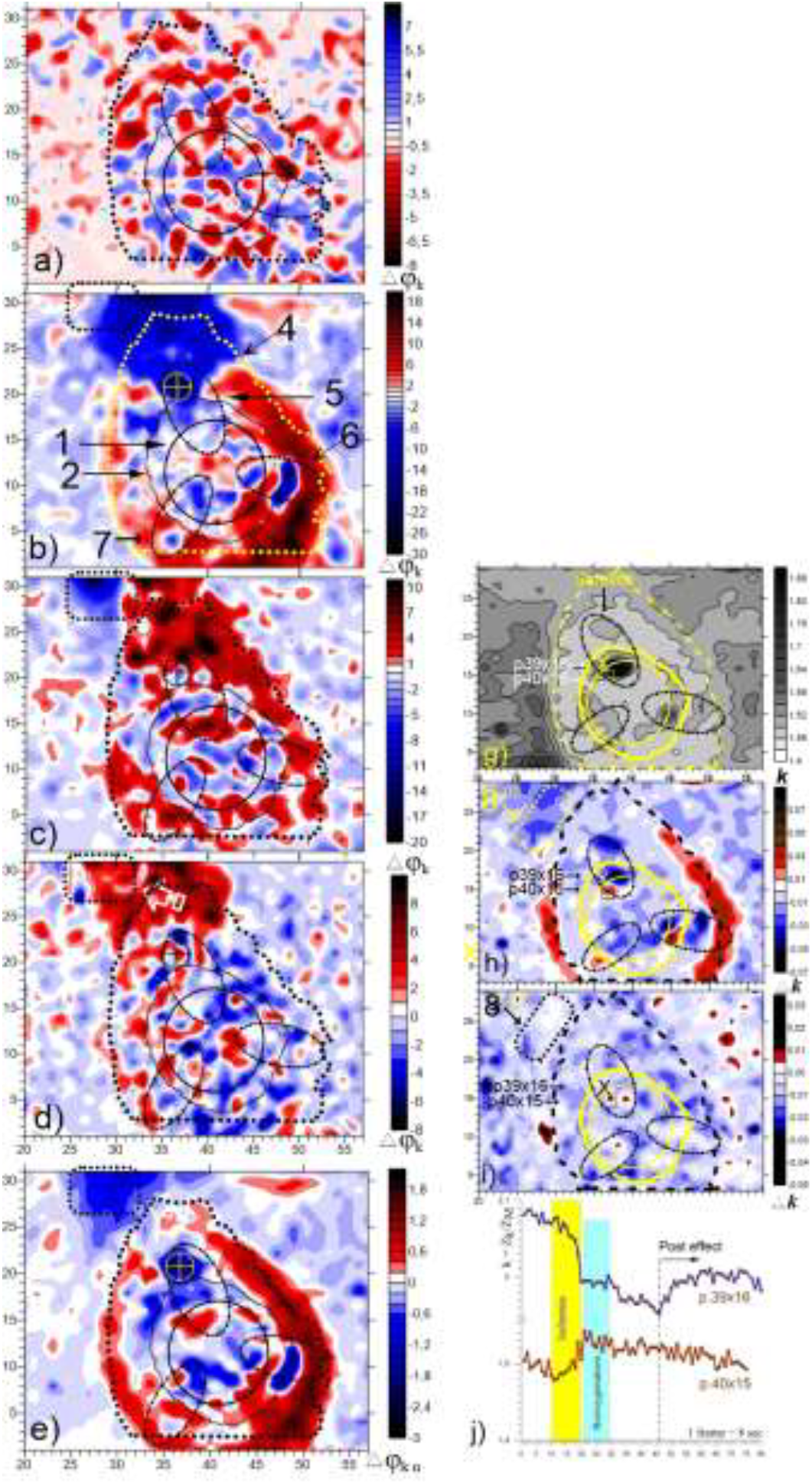
Ischemia-induced dynamics revealing the tumor boundaries. **Left:** The φ_k_-wave generation and propagation in the intermediate zone. The difference normalized maps of the first 4 frames (temporal resolution 8s) demonstrate weak but noticeable initial dynamics of the MS and its surroundings followed by high-amplitude clustering, p<0.01. **Right:** The ion balance maps ***k*** in statics and dynamics (**h**,**i**); **j)** The ***k-*** time curves of two points located on opposite sides of the PN boundary. Designations - see Fig. 2.

**Fig. 5.**
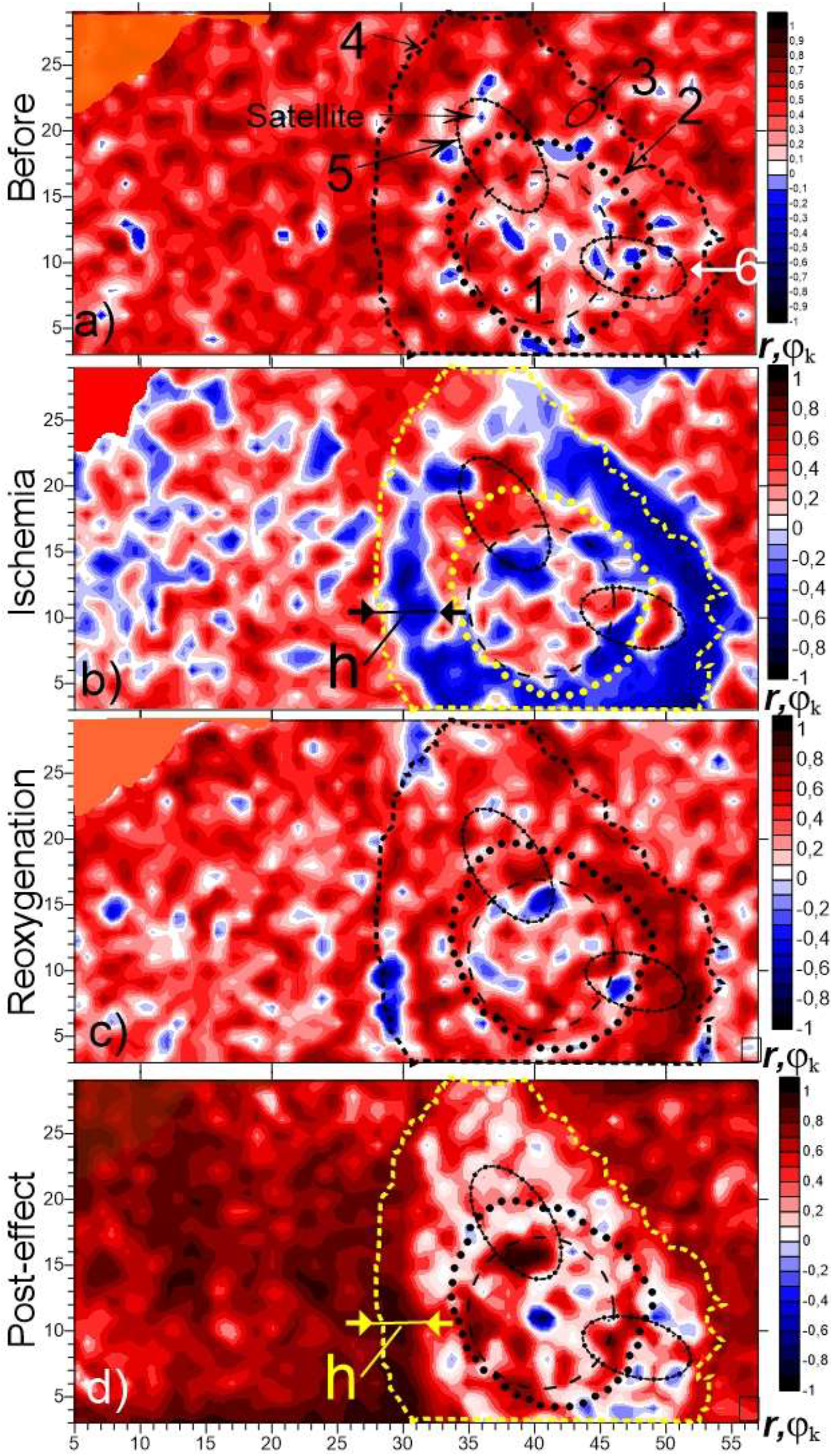
Coherent dynamics of φ_k-l_andscape during the whole experiment. Correlation fields relative to averages reveal (p<0.01): **a)** Initial presence of several remote microclusters and a larger number/size in the “4” boundaries; **b**) sharp antiphase structuring of “3” (−1.0 ≤ r ≤1.0), with a basically important exit beyond “4”; **c**) Relatively rapid relaxation process with disappearance of most antiphase clusters; **d**) Establishment of a maximum stabilization mode during the post-effect period with the disappearance of antiphase clusters everywhere except “4”. Conversely, “3” stand out with even less variability than before.

- Contour of the pre-existing nevus 1 (Fig. 3);
- Unexpected antiphase structure in 6, generally similar to the structure of satellite zone 5, that is, presumably another developing, but not yet visible satellite.

Both of these zones 5 and 6 (as supposed emitters of negentropy) determine the asymmetry of the initial/conditional impedance boundary* 4 (Fig. 3a), as well as the wave-like dynamics in the IZ (Fig. 4).

*Contour of φ_k_ as the most contrasted, Fig.3.

### 1.1. “From chaos to order and back”

Fig. 3a) depict initial SELs in four extra- and intracellular parameters: Z_k_, Z_M_, φ_k_, φ_M._ Fig. 3b-d) illustrates the transition of these patterns in a manner”order from chaos and back” as the fluctuations maps calculated for the initial (b), ischemic (c) and reoxygenation (d) periods (of 15 frames, i.e. 2.5 min each). In the snapshots (Fig. 3a), most of the above mentioned areas of interest are better distinguished in the φ_M-_landscape a4).

As compared with initial stage, the ischemia led to a sharp contrast of the three main zone of interest (3,5,6, Fig. 2) with an increase/decrease in σ by 2–5 times. Moreover, it is worth noting new, previously unnoticed, phenomena:

- A distinct manifestation of IZ as a sharp - almost to zero-weakening of fluctuations* at the level of ECF and ICF (Fig. 3 c1-c2); And, what is especially important in the context of FBT:
- **Expansion of active zones 5**,**6 (i.e.invasive fronts/negentropy emitters) beyond the impedance boundary 4, which was especially evident everywhere for the satellite zone 5 and for zone 6 in the intracellular environment - in contrast to the surrounding weakening 4;**
- Activity in the elliptical zones (both in liquids and membranes) stands out sharply, particularly marked in the intracellular level (c2);
- Increase of the cell and intracellular membranes impedance fluctuations in IZ.

* Similar effects were also registered in response to the weak MF exposure, Fig.S5.

***Reference note***. *The transition from initially chaotic state to more organized patterns and back, when the external impact stops, emphasizes the dynamic nature of biological systems and their ability to self-organize—a key principle in the theory of DS. The “order from chaos” concept suggests that while ischemic conditions introduce chaos at the cellular level, they can also lead to emergent properties where certain groups of cells organize their responses effectively— forming coherent structures that enhance survival strategies or adaptive mechanisms. However, it is essential to recognize that this order does not imply uniformity; rather, it highlights how some degree of chaos (fluctuations) is necessary for adaptability and resilience within tumor. microenvironments*^*36*^

### 1.2. Ischemia-induced dynamics revealing the tumor boundaries

Main events:

- Wave of polarization/depolarization of cell membranes of the intermediate zone, overlapping the FTB;
- Anti-phase dynamics at the invasive fronts;
- Transit change in ionic balance k= Z_k_/Z_M_ at the morphological and functional boundaries of the tumor.

The initial dynamics of the φ_k_–landscape (Fig. 4a) is presented as a difference image between 2d and 1^st^ frames. The color scale is reduced by 2–3 times to show that ***initially the entire region limited by FBT stands out against the background of increased activity in the form of antiphase clusters of small amplitude***. At the same time, it is necessary to note its weak structuring: ***one can only notice a chain of negative clusters around the VB***.

The next difference image ***b*** (between 3d and 2d frames) demonstrates the first response (Fig. 4b), where one can see the ***emergence of two large high-amplitude antiphase structures significantly expanding the FBT 4, and also penetrating inside the VB 2***. It is clearly visible that the epicenters of these structures are three zones: satellite 5 (blue) and two ellipses 6 and 7, as presumable zones of younger invasive fronts. Moreover, in all these three zones, the ***manifestation of smaller double antiphase structures is observed at the borders of the visible tumor 2 and its preexisting nevus (PN) 1***.

The next (4th-3rd) difference image ***c***, compared to the previous one, shows propagation of the red structures (the velocity is about 1-2 mm/sec) with absorption of the blue one. This continues on the next frame ***d*** (5^th^ -4^th^).

This transition was mainly complete by 1 min. The (8^th^ – 1^st^) image ***e*** shows resulting change in the φ_k_–landscape, which then remains almost unchanged. Its further modifications in the re-oxygenation stage are presented as correlation fields in Fig.5 and as fluctuation maps in Fig.3c3. As it follows from the patterns of Fig, 3c, a topologically similar response to ischemia can be in more or less degree traced on the all SEL parameters.

As it was firstly reported^34^, the functional heterogeneity of the entire melanoma zone can be estimated in clinic by a single snapshot of the SEL, i.e. as a distribution of the dimensionless parameter of ion balance k = Z_k_/Z_M_. Fig. 3g shows such a distribution, as well as its change at the stage of ischemia and reoxygenation, Fig. 3h,i. On the map of Fig. 3g, all three zones outlined by ellipses stand out clearly, as well as the boundaries of the quasi-stationary FBT. However, more informative are the difference of ***k-***maps for the periods of ischemia and reoxygenation (Fig. 3h,i), they:

- contrast the zones of increased reaction to ischemia both inside the FBT and outside -8, and also show the reversible nature of these changes;
- demonstrate the phenomenon of the FBT expansion (red structures) by 5-10% increase of ***k***, Fig.3h.

Fig.3j gives an idea of the antiphase of the time dynamics of ion balance points located on different sides of theVB. Moreover, this applies to both the ischemia period and the post-effect stage*. It is noteworthy that ***the process of changing the ion balance in anomalous zones exhibits inertia\*\**** compared to the time dynamics of the parameters Z_k_ and Z_M_ separately, which is consistent with known indirect evidence that the temporal dynamics of ECF and ICF conductivity in cancer-affected tissues is likely to be asynchronous and slower compared to healthy tissue^60^.

* The latent period approximately coincides with that in damaging a plant leaf (Fig. S16).

** The conductivity/impedance of a medium is directly related to the concentration and mobility of charge carriers. Within the ICF, the primary charge carriers are ions such as K+, Na+, Cl-, and various organic anions. In the, Na+ and Cl- are dominant. Changes in the permeability of the cell membrane due to altered ion channel activity will affect the rate at which ions move across the membrane, thereby influencing the temporal dynamics of ion concentrations in both compartments. If ion channels or their gating properties are altered as in the tumor microenvironment or FBT, the rate at which the ECF and ICF ionic environment responds to a stimulus is also altered. The absorption of the blue structure by the red one (Fig. 6c) can be associated with cooperative depolarization and subsequent hyperpolarization of cell membranes.

**Fig. 6.**
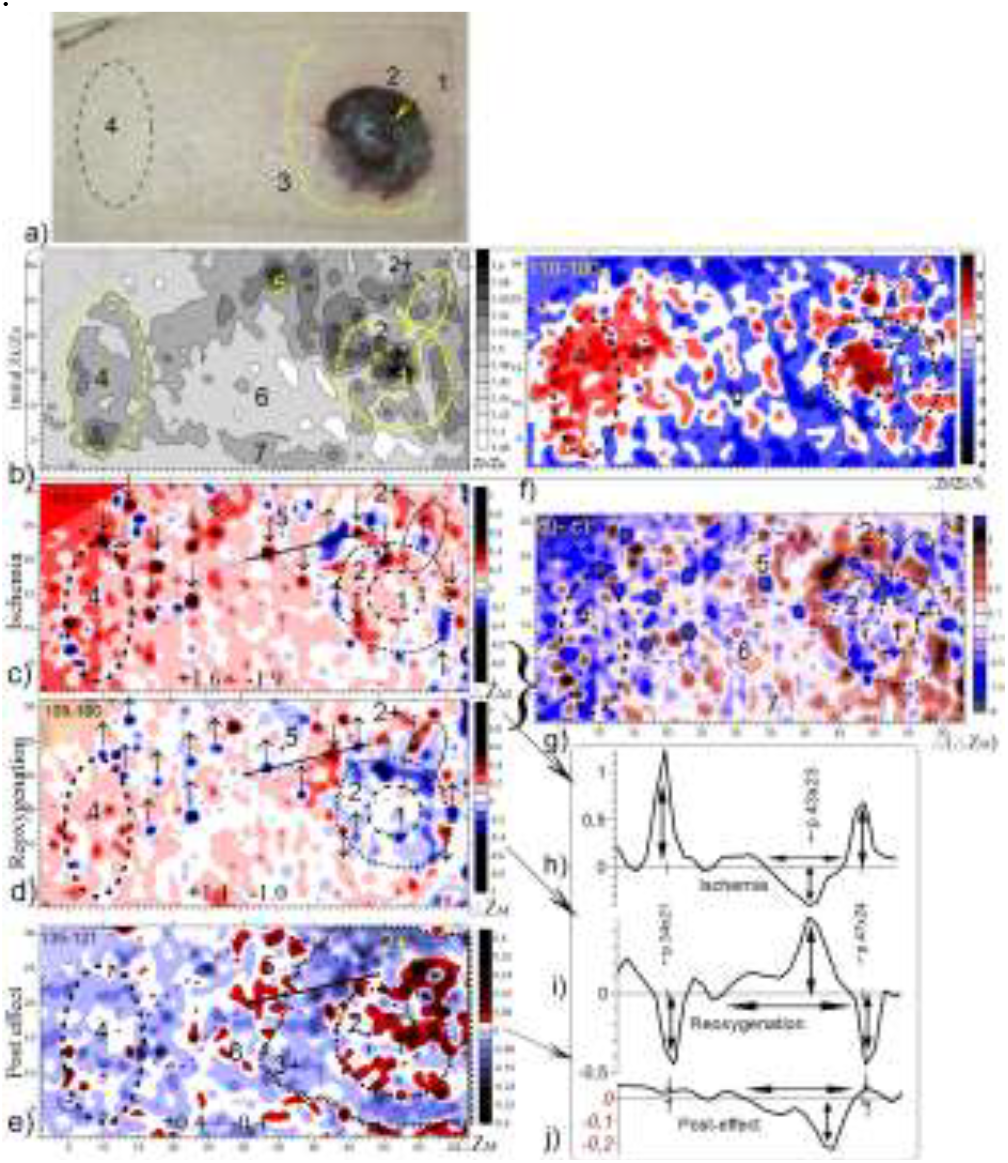
Emergence and Expansion of Functional Boundaries at the Level of Cellular Liquids. **a)** photo of SSM. 1-PN, 2-visible boundary of SSM, 3-boundary of φ_M_ landscape (see Fig.7b), 4,5,7-zones of abnormal IB k=Zk/Z_M_, 6-intermediate region; **b)** calculated map* of k; **c)** Difference of normalized Z_M_ images for the ischemia period (frames 100-91, i.e. ab.1.5min). Arrows ↓↑ indicate peak points with antiphase dynamics **d)** Similarly, the reverse changes of all marked peak points during reoxygenation (frames 109-100); **e)** Difference map for the post-reoxygenation period (frames135-121). Line 3+ denotes the advance of the front 3; **f)** map of k changes during ischemia; **g)** Difference map of (**d**) and (**c**); **h-j**) Profiles drawn through the peak points 34×21 and 47×24 (marked with a line at **c** and **d**), p<0.001.

***Reference notes***. *The observation of autowave processes and antiphase structuring in biotissues serves as compelling evidence for their dissipative origins, demonstrating that biological systems can spontaneously organize into complex patterns when subjected to weak stimuli, consistent with the theoretical framework of DS. The phenomenon illustrates the dissipative nature of the melanoma zone through the interaction of various cellular ensembles in response to ischemia. The most aggressive boundaries/zones of the tumor, are often the most metabolically active and thus can be defined as main emitters of negentropia*^*36*^. *The activation of younger cellular ensembles in response to ischemia highlights a fundamental characteristic of DS: they maintain order and structure by exchanging energy/information and matter with their surroundings. This is especially obvious as the zone 6, presumed to be more active, responds more vigorously to short ischemic events compared to its visible counterpart. As ischemia induces changes in the impedance landscape of cell membranes, coherent structures emerge on either side of the VB. The oppositely directed coherent structures signify a dynamic equilibrium that is characteristic of DSs where energy dissipation leads to self-organization*.

### 1.3. Ordering and disordering phenomena in φ_k_-correlation fields during ischemia and reoxygenation

Fig. 5a-d depict the relative averages of the φ_k_-correlation fields for each. Main features:

At the initial stage (**a**), the melanoma area is somewhat distinguished by an increased level of spatial synchronization outside the tumor, going beyond the limits designated by FBT 4, against the background of a reduced level - up to islands/clusters - of negative ***r*** values - in the zones of invasive fronts 5.6;

The reaction to the ischemia stage (b) can be defined in many ways as the *opposite of the initial dynamics* of the entire zone, especially of the melanoma region:

- antiphase dynamics (−1.0 ≤ r) of the intermediate zone with a significant exit beyond the FBT. At the same time, zones 5, 6 are distinguished by the opposite dynamics (r ≤1.0);
- negative clusters appear in the distance.

The reoxygenation stage is characterized by reverse dynamics in the form of leveling of distant negative areas and a distinct reversal of the intermediate zone 3 (r ≤1.0).

***Of particular interest is the systemic response revealed as a post-effect stage (d) in the form of counter-directional ordering of intermediate zone “3 “and its surroundings, as presumably normal tissue, i*.*e*.:**

- ***Disordering of the intermediate zone demonstrating the noticeable “retreat of the front-line” as a contraction of its functional boundary;***
- ***Ordering spatial correlations of the surroundings up to r≤1*.*0 with the relative “advance of the front-line”***, *max h≈3mm*.

***Reference notes***. *The host system’s response to ischemia is likely intended to restore order and homeostasis. The ordered response in the melanoma macroenvironment could be the host’s attempt to control the tumor. When the tumor zone becomes disorganized, it could be interpreted as the tumor’s ability to withstand the stress of ischemia. The host system might be attempting to disrupt the tumor’s structure and function. However, the tumor’s response (disorganization) could be a mechanism to survive the stress. This could involve the tumor cells activating survival pathways, altering their metabolism, or changing their interactions with the surrounding environment*.

#### <u>Analysis II</u>. Superficial Spreading Melanoma^33^ (SSM). Energy and information impact

The plan of the entire experiment is described in [33] and is also clear from Fig. 9. In contrast to MS, this case is notable for the absence of visible abnormalities (Fig. 6a), but the presence of invisible large abnormalities detected in the initial quasi-stationary SEL, which could presumably mean correspondingly extended functional tumor boundary, which *may exceed the entire scan-area*. This boundary is designated as general FTB. In contrast, we will consider line ‘3’ as the nearest “front line” defined by the clearly visible outer contour of the mitochondrial* landscape, Fig. 7b.

**Fig. 7.**
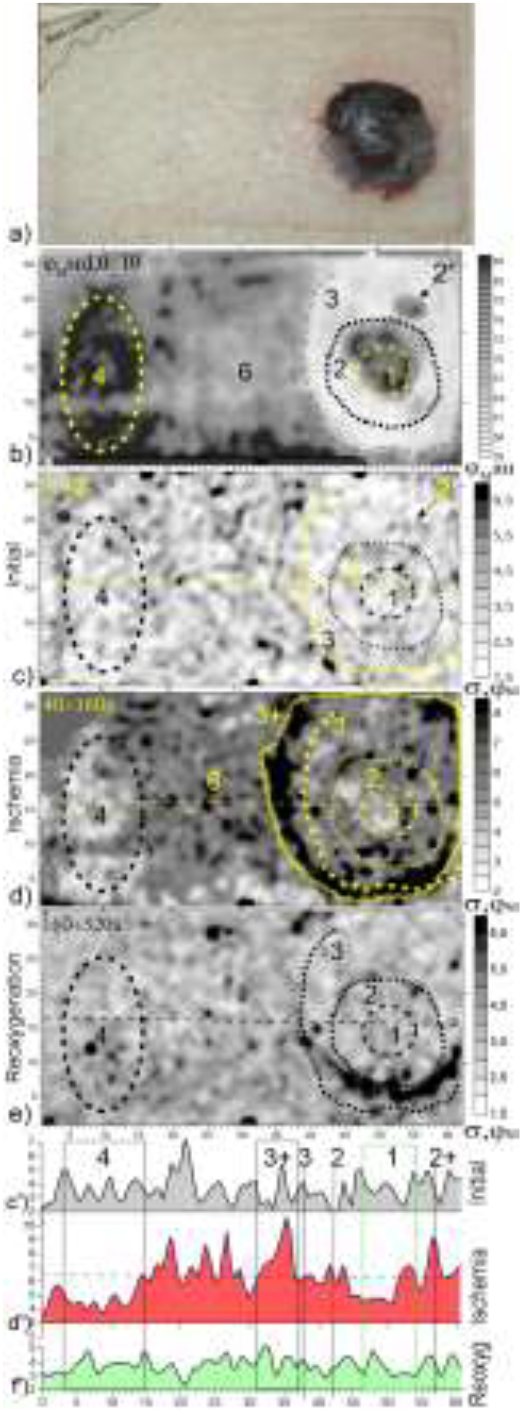
Radial expansion of mitochondrial functional boundaries in the fluctuation field during ischemia. **a)** Photo; **b)** Initial φ_M_-landscape; .**c)** Initial dispersion field (the scale has been increased); d) Increased fluctuations and expansion of the boundaries to “3+” in the response to ischemia; **e)** Weakening and smoothing of the fluctuation landscape; **f)** Landscape profiles (dotted lines in the middle of the c,d,e frames), p<0.01

**Fig. 8.**
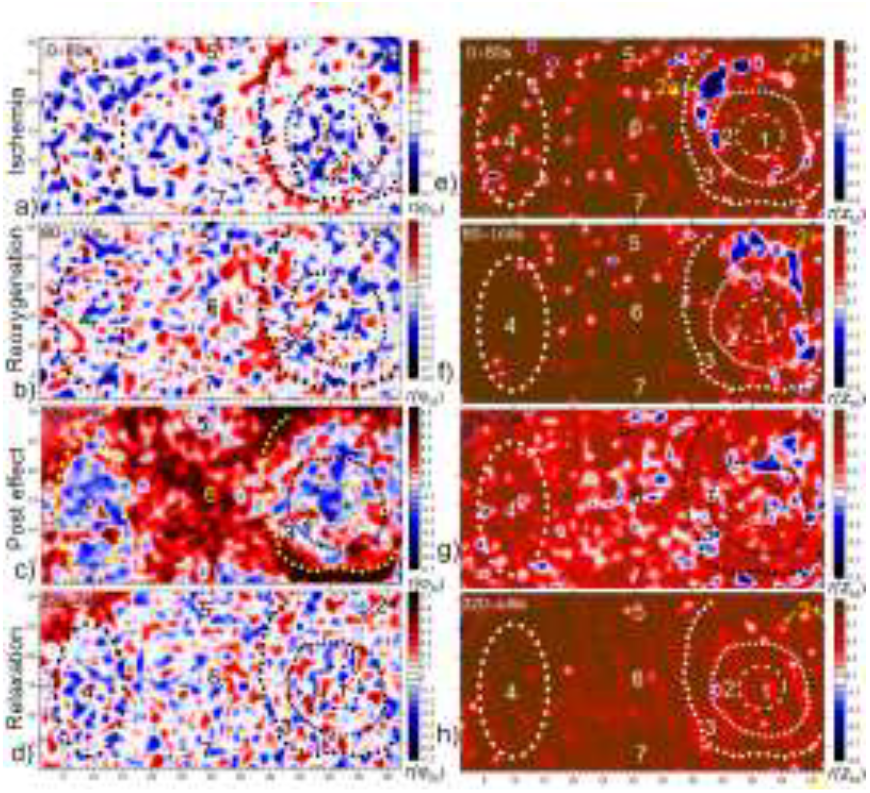
Differences in the dynamics of the mitochondrial membrane landscape (left) and intracellular fluids (right) throughout the experiment. Correlation fields of φ_M_ and Z_M_ (**e-h**) for all stages. Of note: **Left**: Emergence of a red structure on the mainly chaotic landscape-clearly along line “3” (**a**); its predominant expansion to the left and emergence of a red structure along line “4” (**b**); further spread of red into “6”, skirting zones “3”, “4”, “5”, “7”, with emergence of zones of antiphase dynamics in the latter (**c**); relaxation to the chaotic regime (**d**). **Right:** Emergence of point clusters and the “2a” structure (blue) on a highly stable landscape (**e**); continued emergence of point clusters, *disappearance of “2a” and appearance of “2+”* on a still highly stable landscape (**f**); general decrease in orderliness up to the appearance of negative clusters mainly in (6), and with noticeable dynamics of “2+” (**g**); stabilization of the landscape with disappearance of point clusters (**h**).

**Fig. 9.**
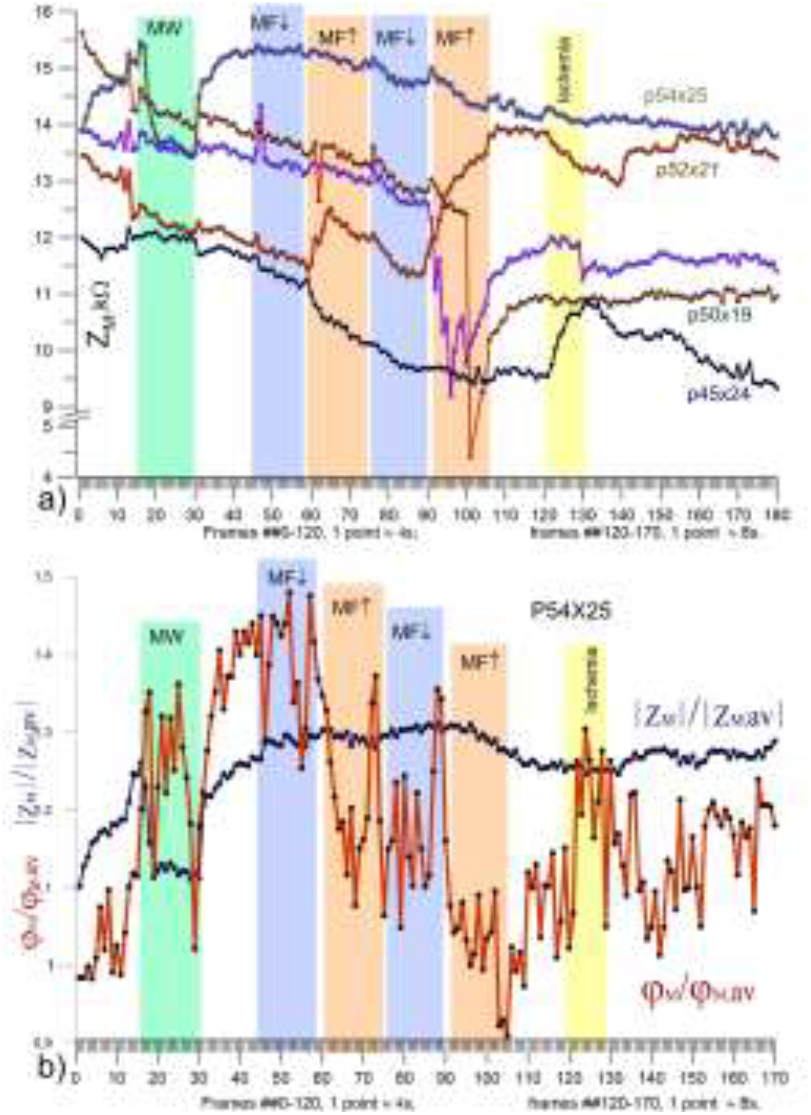
Diversity of temporal response of individual points of the Z_k_ and φ_M_ landscape throughout the whole experiment. **a)** The Z_M_ time courses of five randomly selected pixels; **b)** A point* of the 2+ area demonstrates the high sensitivity of membrane impedance compared to liquid impedance (as relative to the averages), p<0.001. *In coordinates 64×32

*Since the mitochondrial membrane is a key target in ischemia, and there’s some evidence suggesting potential effects from nonthermal EMF, particularly in cancer cells^46,49,50^.

As in MS, the dimensions of IB of MSS significantly exceed the apparent size of the tumor (indicating real FPTs), and they, as above, do not have clear boundaries. However, the ion balance map k = Z_k_/Z_M_ clearly reveals the presence of an extended anomaly “5” exceeding the background by ∼10%, (Fig. 6b, Supplementary Materials I, Fig. S1). It is noteworthy that this anomaly comes directly from the PN and, stretching in two arms* across the entire frame, ends with the suspicious structure “4”.

Another reason for choosing the SSM data for this more detailed analysis was possibility to compare effects of both factors: energy (ischemia) and informational one (MF/EMF) used in this experiment. In both experiments similar ischemic test protocol was used, i.e. stopping blood circulation in the limb with a cuff for 1.5 minutes.

*Sleeve “7” starts from the VB, probably due to its later origin.

### 2.1. Energy/Ischemia Impact

#### 2.1.1. Emergence and Expansion of Functional Boundaries at the Level of Cellular Liquids. What could be hidden behind the Landscape of Ion Imbalance?

Fig. 6 demonstrates the ICF spatial collective dynamics Z_M_ (as more susceptible to ischemia than Z_k_) and the resulting changes of k = Z_k_/Z_M_ 2D response to the transit ischemia.

The difference image of Z_M_ landscape for the ischemia stage(Fig. 6c) showed that in the region “5”, in phase with the background, a family of peak points* (microclusters) appeared, as well as a pair of intratumor points (in “2”), marked ↓. At the same time, a family of out-boundary points marked ↑ showed opposite/antiphase dynamics.

*\*Presumably hypoxic niches, microsatellites or in-transit metastases*.

The stage of reoxygenation reveals:

- Amplitude reversal;
- **Expansion of the area of these boundary clusters** (red). This is also shows graphically in Fig.6i,j.
- Similar but oppositely directed process occurs on the inner side of “2”, resulting in the **formation of an antiphase structure on both sides of the VB 2**.

The post-effect manifests itself in an even larger-scale ordering of the Z_M_–landscape (Fig. 6e,j):

- **Coherent**/**in-phase expansion of the near surroundins to the “front line” “3+”(blue), with the appearance of antiphase (red) clusters on the outer side of “3+”, forming thus as a whole a new antiphase structure-now at the border of “3+”**. At the same time, the structure of “4” remains almost indistinguishable (compare with Fig. 8c);
  - Clustering of PN zone “1” with increasing contrast of zone “2+” (red);
  - Smoothing of the entire landscape, especially to the left of “3+”, with the disappearance of point anomalies, which is clearly visible when comparing profiles p.34×21 with p.47×24, Fig. 6j.

The resulting change in the ***k***-landscape is shown in Fig. 6f as a difference image over the period of ischemia. As expected, it highlights: zone PN and the immediate vicinity of the tumor “2” (including zone “2+”, Fig. 7), as well as - unexpectedly - **similarity between zones “4, 4+” and the tumor zone** (compare with Fig.8c).

#### 2.1.2. Emergence and Expansion of the Mitochondrial Functional Boundaries. Asynchrony of Spatiotemporal Dynamics of Mitochondria and ICF

The initial snapshot of φ_M_-landscape and its dynamics are depicted in Figures 7 and 8.

Fluctuation field of the preliminary stage (Fig. 7c), despite the zoomed-in scale, is a featureless chaotic landscape (σ_max_=12).

The ischemic stage (Fig. 7d) demonstrates **emergence of a new ordered structure “3_3+”** (σ_max_ =48) at about totally increased fluctuation landscape. Of meaningful exception, characterizing their similarity, are two areas: **“1” and “4”, they are equally manifested as** an unchanged level of pulsations, which is also shown in the relief of Fig. 7d’.

At the reoxygenation stage (Fig. 7e), one can see a significant and ubiquitous weakening (σ_max_=27) and smoothing of the fluctuation landscape compared to the original (c).

Fig. 8 demonstrates the differences in the spatiotemporal dynamics of mitochondria and ICF in the form of their correlation fields of their impedance throughout the whole experiment. ***The collective dynamics of mitochondrial membranes and ICFs reflect different/ complementary informational signatures: they are not synchronized and show different responses***.

In general, the correlation field of ICF looks much more stable than that of mitochondrial membranes due to the strict ionic homeostasis of the ICF, while mitochondrial membranes are subject to various factors that can affect their stability^61^.

The main feature of the mitochondrial scenario is the emergence of a coherent structure on the initial impedance boundary “3” during ischemia (Fig.8a) and its subsequent wave-like expansion* in such a way that all suspected zones, especially “4”, become contrasting by their antiphase dynamics – the same as in the tumor, Fig. 8c.

* *In more details, this process is presented in Fig.S2 (Suppl. Mat. I)*.

The ICF dynamics does not reveal the mitochondrial phenomena of expansion of “3” and contrasting of zones “2”&”4”. Here, instead, against the background of a stable landscape, two large structures* within “3” clearly appear one after another: “2a” and “2+”, which are less distinguishable on the mitochondrial landscape. Also, in contrast to the latter, at the stage of ischemia and reoxygenation, the antiphase microclusters (see Fig. 6 c, d) are clearly distinguished here, which disappear at the relaxation stage. The systemic reaction in the post-effect stage manifested itself as a general decrease in orderliness up to the appearance of negative clusters mainly in (6), and with noticeable dynamics* of “2+”.

*\* The same areas were highlighted in response to EMF exposure, see also Figs.9, 10, 11e-h*.

**Fig. 10.**
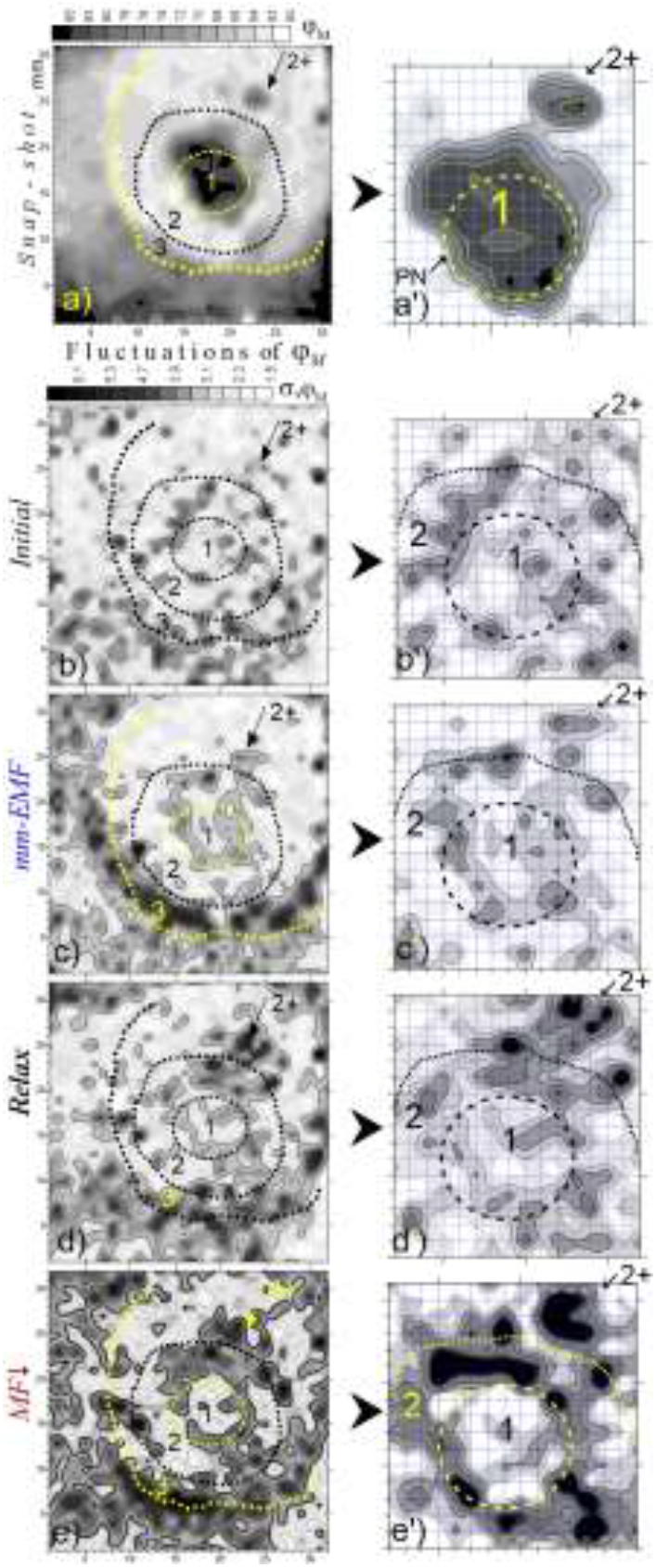
The mm-EMF/MF_↓_-induced φ_M_ fluctuations. **Top row)** The φ_M_ and Z_M_ snap-shots; **b-e, g-j)** their fluctuation maps; **a’-e’)** φ_M_ in an enlarged scale; p<0.001.

**Fig. 11.**
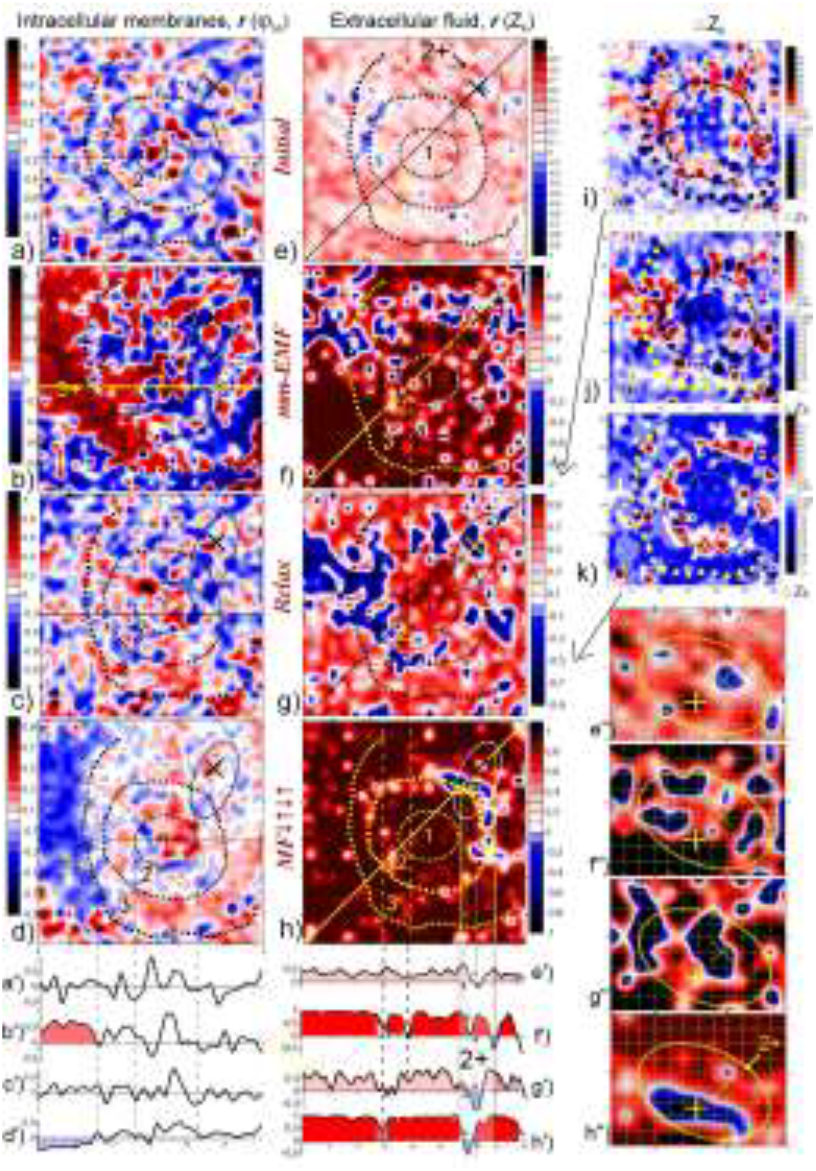
Effects of clustering/declustering and DS movement. The φ_M_ and Z_k_ correlation fields relative to average: **a**,**e)** Initial stages**; b**,**f**) The mm-EMF stage; **c**,**g**) Relaxation stage: **d**,**h)** The MF exposures (see Fig.8). **Bottom**) Profiles of a…h .(solid lines) **Right column**) The DS effect of “reversible structuring”: **i-k)** Difference Z_k_ maps during the relaxation period (**g**); **e’’**..**h’’**) Enlarged fragment of **e_h** showing antiphase dynamics of supposedly true boundary points marked as “×” and “+” in (h). p<0.01.

In summary, the post-ischemic response can be characterized as either ordering or disordering based on the observed patterns of correlation fields. The presence of contrasting patterns suggests ordering, while the disruption of landscape stability indicates disordering. Within the framework of self-organization theory, the **antiphase correlation in the tumor surroundings – the red zone in Fig.8c - can be a sign of normal tissue response\***, but further investigation is needed. While mitochondrial behavior is a valuable indicator of tissue state, “normality” cannot be solely defined by mitochondrial function. A holistic approach is necessary to understand the complex interplay of cellular components and processes in the context of tumor and its environment.

* ***This assumption may also be supported by the similarity with the above-presented MS case, i*.*e. with the effect of “advancing the front line” of the tumor surroundings*** *(compare Fig. 5h and Fig. 8c)*.

### 2.2. Information impact: EMF/MF-induced reorganization of SEL

The assumption that EMF and MF has a more specific effect on the tumor macroenvironment compared to ischemia is grounded in the idea that EMF/MF can modulate cellular communication and metabolic processes in ways that ischemic conditions cannot replicate.

The entire course of the experiment described in [33] and can be seen in Fig. 9a, in which the mm-EMF/MF part is presented in 7 stages of 1 min each. (A 10-min break was made between the EMF/MF test and the ischemic one.)

To double the temporal resolution up to 4 s/frame, the SSM was read in 1 half of the scanning head (32×32mm^2^).

#### 2.2.1. Asynchrony of some Representative Points

##### Locally increased sensitivity of φ_M_ compared to Z_M_

Fig. 9a shows the time curves of five pixels inside and outside the contour “2”, thus demonstrating different scenarios, noticeable differences in response time, and large differences in the sensitivity of the cytosol impedance Z_M_ to each of the applied stimuli throughout the experiment. In principle, for each of these points, as for all the others, it is possible to conduct a correlation analysis to identify the coherence zone surrounding it. However (since such voluminous work is best left to AI), we had to limit ourselves by spatial fluctuations of Z_M_ and φ_M_ in response to mm-EMF and their average response to all four stages of MF.

Peculiarities of the zone “2+”, in particular, manifested themselves in the initial dynamics of the pixel p. 54×25: the antiphase dynamics of Z_M_ (Fig.9a) and the high amplitude fluctuations of φ_M_ (Fig.9b). Many other points, especially at the mitochondrial level, with abnormal initial dynamics are seen in the fluctuation maps, Fig.11b,g.

Interestingly that Z_M_ in “2+” (p. 54×25) exhibits high sensitivity, unlike that of φ_M_, to only one factor – mm-EMF, Fig.9a.

*Note. In the context of the ischemic test results, it is worth emphasizing that the membrane response (φ*_*M*_*) of some specific regions was also significantly more marked than that of the fluid resistance (Z*_*k*_,*Z*_*M*_*), which is especially noticeable, for example, in the intriguing “2+” zone, as shown in Fig. 9b*.

#### 2.3.2. Mitochondrial Fluctuation Field: Expansion of FTB and Emergence of Stroma of Preexisting Nevus

Fig. 10 presents fluctuation maps of φ_M_ for the four first stages of the EMF/MF experiment. (The entire set of φ_M_ fluctuation maps is shown in Fig. s4). Fig.10b depicts initial landscape, that is more chaotic than not; Fig. 10c) shows that the impact of mm-EMF leads to a sharp ordering of the fluctuations landscape and manifestation of:

- associated structures in zone “1” (presumably stroma of the PN) and 2+;
- a girdle structure along line 3 – the initial quasi-”stationary” boundary of the mitochondrial landscape in real (non-fluctuating) parameters.

With mm-EMF switched off, collapse of these structures begins (Fig.10d), but subsequent exposure to MF↓ (Fig.10e) restores the ordering with an even greater amplitude of fluctuations.

#### 2.3.3. Correlation Fields of φ_M_ and Z_k_: Ordering-Disordering Phenomena

- *Emergence and reversal of the mitochondrial front in response to mm-EMF/MF;*
- *Dual ordering of Z*_*k*_ *-correlation field in response to EMF and MF, characterized by predominance of high positive correlation in the background, juxtaposed with isolated regions/clusters of negative correlation (supposed DS) at the peritumor area;*
- *Direct evidense of DS: reverse migration of Z*_*k*_ *clusters to the tumor border*.

Fig. 11 shows differences in the processes of ordering the φ_M_ and Z_k_ landscapes throughout the entire EMF/MF experiment in the form of correlation fields.

The most significant features of the φ_M_ and Z_k_ dynamics are:

**1) (a**,**e**) Both landscapes reflect predominantly chaotic dynamics, where individual coincidences with the tumor structure are probably explained by the influence of measuring microcurrents^51^.

In both cases, similarities were observed:

**(a)** large-scale structuring of φ_M_ field throughout the field and significantly increased clusterization in the PN “1” (see also the **a’** relief);

**(e)** small-scale structures Z_k_, particularly noticeable in the “2+” zone and in the left upper peritumoral zone, i.e. in the same peritumoral area where they appeared in response to ischemia (Fig.5).

**2) (b**,**f**) Both landscapes reflect the process of sharp ordering in the response to mm-EMF. At the same time, in (b), as in the response to ischemia (Fig.6d), the *same phenomenon of emergence of the coherent “3_3+” structure of φ*_*M*_ is evident. The response of the Zk (f) changed towards greater polarization of the whole landscape thus highlighting areas with antiphase dynamics against the background of the majority with r≤+1.0. This impressive phenomenon of ordering is clearly visible in the profiles. (e’-h’). (Notably, the topology of negative clusters basically corresponds to that under hypoxia Fig. 6c).

**3) (c**,**g)** At the relaxation stage, one can see the reverse dynamics, i.e. disordering of the field φ_M_ and decrease in the coherence of the red background of Z_k_ (g’) with amazing simultaneous growth of the blue structures, as a continuation/aftereffect of the response to mm-EMF. It should be borne in mind here that the correlation field was calculated based on 15 frames of each series and thus represents an averaged estimate of this dynamics.

To reveal it in more details, difference images of the Z_k_ for this period were calculated. Fig. 11 i..k show that this process occurs in the form of *contraction of the family of dissipative structures* (blue) to the VB “2”. (Without detecting this dynamics, the following changes. i.e. in ***h***, could be erroneously attributed to the action of the MF). In Fig.11k, the original feature of the “2+” zone as the end point of this “contraction” (and, hypothetically, a separate emitter of negentropy) appeared in the form of a separate DS marked with “×”.

4) (**d**,**h**) During the MF stage, high ordering of Z_k_ (r ≤+1.0) manifested itself almost on the entire landscape “h”. From the “blue” pattern basically only “2+” zone with a few neighbouring clusters at the tumor border remained. To emphasize the antiphase nature of two adjacent points this zone, the figures (e’’-h’’) present the process of the “2+” rearrangements on a larger scale. These points (“×”, “+”) are probably located on opposite sides of the true morphological border of the tumor (in clarification of the approximately designated boundary “2”). Regarding the φ_M_ dynamics, the most significant is the secondary manifestation of the “3_3+” structure as a negative correlation. In addition, a noticeable smoothing of the landscape can be noted (d,d”).

In general, the main similarity of the collective impedance dynamics of mitochondria and cell liquids in response to weak energy and information impact (ischemia, non-thermal EMF) is that in both cases there is an expansion of FBT in the form of activation and propagation of the front dynamics of *corresponding* initial impedance landscape. A significant difference is that, unlike the mitochondrial response, the cellular fluid response is manifested predominantly only on the outer contour of the tumor, Fig. 15.

#### <u>Analysis III.</u> Irradiated Melanoma: Disruption or Weakening of Intercellular Communications Everywhere but Cellular Membranes

Unfortunately, the protocol for this experiment did not include an ischemic stage, but was otherwise similar: mm-EMF and a sequence of oppositely directed MF fields (Fig.9a).

Fig. 12 depicts static and dynamic features of spectral impedance landscapes: k = Z_k_/Z_M_,φ_k_ and φ_M_. Their fluctuation fields ***e-h*** are presented for the most active stage – mm-EMF. The absence of noticeable collective ordering can be seen everywhere but at the level cell membranes ***g***. Unfortunately, the protocol for this experiment did not include an ischemic stage, but was otherwise similar

As in all the presented cases, the static IB noticeably exceed the visual ones. Also, in response to mm-EMF, the φ_k_-field (Fig.6g) demonstrates the phenomenon of additional expansion of these boundaries (Fig.6c).

**Fig. 12.**
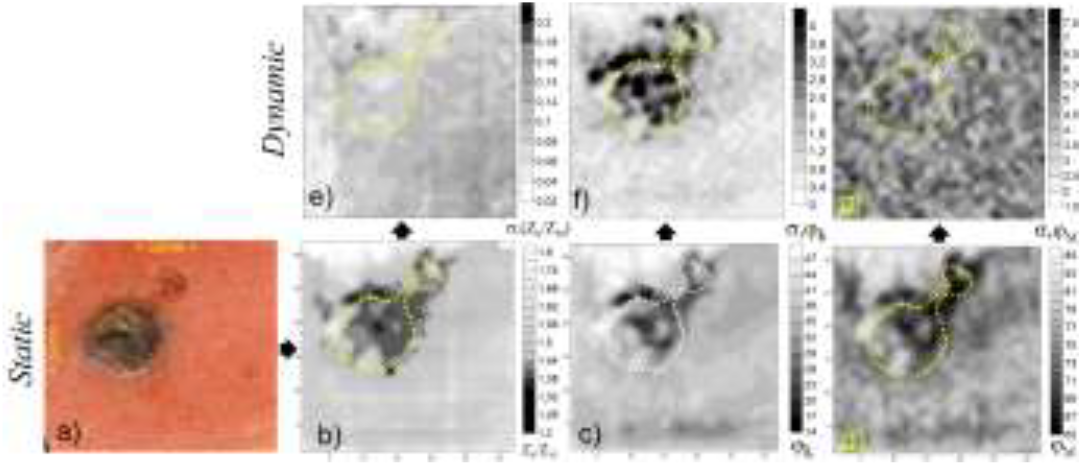
No pronounced response to EMF at all levels except cell membranes. **a)** Photo of the X-ray irradiated melanoma; **b-d)** initial impedance landscapes: ionic balance k= Z_k_/Z_M_ (**b**), cellular φ_k_ (**c)** and intracellular (φ_M_) membranes (**d**); e-g) their corresponding fluctuation fields in the stage of greatest response (mm-EMF).

The behavioural features of membranes at the initial stage are also visible in the correlation field Fig. 13a. Against a background of strong positive correlation (close to 1), the specific tumor zone looks like an area of disrupted correlation. The difference images of φ_k_-landscapes for the same period give the resulting map of changes (Fig. 13b), where the tumor zone shows relatively increased dynamics, which is similar to the dynamics of the non-irradiated MS tumor (Fig. 4b). At the mm-EMF stage (Fig. 13c), one can also note the expansion of these boundaries in the form of large-scale clustering. However, unlike MS, here **the phenomenon of contrasting the boundaries in the form of antiphase clustering is weakened or absent**.

**Fig. 13.**
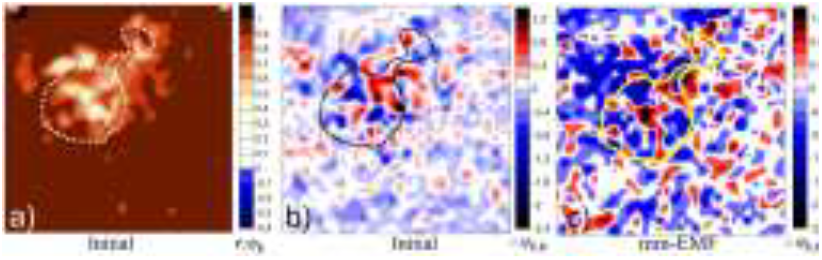
Initial and EMF-induced dynamics at the cell membranes level. **a**,**b)** Correlation and fluctuation fields, respectively; c) Difference image for the mm-EMF stage period

It should also be emphasized that even in the case of ionizing damage, such important common features of all the presented cases are preserved:

- **Excess of the initial IB over the VB**,
- **Additional excess of the φ**_**k**_**-boundaries in the response to mm-EMF due to the survived communications of cell membranes**, Fig. 12g.

#### Summary&Conclusion

The FBT is considered as a notional boundary of a dynamic intermediate zone (IZ)—a “battlefield”—between the hostile ecological system of the tumor and the host, driven by the interplay of cellular heterogeneity and environmental factors. In the presented context, the FBT can be defined as a conditional impedance boundary (IB) beyond which more stable dynamics of the impedance landscape are observed in response to stress, which indicates normal homeostasis, Fig.14.

**Fig. 14.**
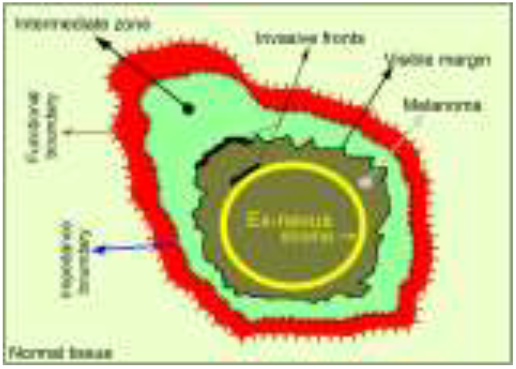
Schematic representation of melanoma and its boundaries. Configuration of the functional boundary is determined by the topology and aggressiveness of the invasive fronts

For the first time – using SEI, it was found that **the initial/static IB may significantly exceed the VB**, what is explained by the sensitivity of impedance to factors such as tumor heterogeneity, microvascular changes, inflammation, ECM remodelling and metabolic activity. These factors can alter the electrical properties of the tissue without necessarily causing significant changes in the optical properties.

Previously^33-35^, we reported fundamentally new SEL phenomena like **wave-like dynamics and antiphase structures emerging in melanoma surroundings in response to short weak stimuli of both energetic and informational nature**: (i) ischemia and (ii) non-thermal mm-EMF and MF. At the same time, **significant spatial and temporal differences in the response of the spectral SELs**, i.e. in the collective processes at the level of intracellular and intercellular membranes and fluids.

From biological perspective, the observed effects of coherent structuring of the SEL are likely the result of a combination of factors, including the increased sensitivity of the tumor-altered tissue and the dysregulated cooperative effects of intercellular interactions within the functionally altered environment of the tumor. This includes altered cell membrane properties, changes in ion channel expression, and modifications to the ECM, all of which can influence the electrical properties of the tissue and its response to external stimuli. It is necessary to emphasize here that, in contrast to the electrodynamics of individual cells, the phenomena presented reflect the *average collective* electrodynamics of multi-thousand cell ensembles underlying self-organization processes.

The intricate biological interpretations of the phenomena, while seemingly complex, find a natural explanation within the established framework of self-organization and dissipative structures. The theoretical lens posits that the observed patterns emerge spontaneously from the interplay of energy/information dissipation and feedback mechanisms, offering a concise and unifying perspective on these phenomena. This approach simplifies the understanding of complex biological systems by focusing on the underlying principles of self-organization. This perspective allows for a more straightforward understanding of the observed patterns. Moreover, such interpretation seems more adequate because it accounts for the complexity of the tumor environment, the role of weak stimuli, and the emergence of ordered structures from disordered states.

Here we presented the results of detailed analysis and synthesis of the datasets from the 4 in vivo experiments, as well as -1 in vitro experiment (plant tissue) as additional evidence for the broad spectrum of the antiphase structuring phenomenon. This phenomenon was clearly manifested mainly at functional and morphological boundaries and, presumably, can also be considered an indicator of tumor invasive fronts.

From the point of view of the DS theory, the observed shape of the IB may reflect the configuration of the total front of entropy flow. The latter corresponds to the topology and activity of local invasive fronts (including e.g. that of ex-nevus), not to the VB shape. This was especially clearly manifested in the shape of the IF greatest expansion at the nevus zone as a result of the increase in the entropy flow caused by ischemic stress.(Transient ischemia further exacerbates its hypoxia, leading to increased metabolic stress and the accumulation of damaging byproducts^88^, thus increasing the entropy flow).

The fluctuation and correlation fields methods, used to analyse induced SEL dynamics, revealed a number of new spatial dynamic features of the functional heterogeneity inside and outside VB, most of which can be understood as how order can emerge from seemingly chaotic landscape - the fundamental concept of the DS theory.

Tumor cells and tumor-affected cells in the IZ often exhibit a lower threshold for responding to stimuli compared to normal cells. At the same time, tumor-affected cells may display a heightened sensitivity to certain stimuli compared to the core tumor cells due to their unique microenvironment and the early stages of transformation they are undergoing^60^. This means that the weak stimuli that would not affect normal cells can trigger the more significant changes in IZ. In this sense, given the demonstrated high sensitivity of IZ, it can be defined as a pre-bifurcation zone. Moreover, the presented results also demonstrated the significant spatial and temporal differences in the IZ assessed at the intercellular and intracellular levels.

A number of similar effects and phenomena of self-organization and dissipative structuring have been registered not only in the area of such a malignant tumor as melanoma, but also in outwardly similar skin anomalies, such as a papillomatous nevus. These main common and complimentary features are presented in the Supplement Materials II.

The abundance of the presented phenomena is explained by the “discoverer effect” of a previously unknown object – the electrical impedance landscape of the skin as such and its spatial and temporal dynamics. However, despite the promising nature of these findings, we acknowledge that the limited sample size and inherent heterogeneity among the studied objects pose challenges for comprehensive biological interpretation. The variability in tumor characteristics may lead to difficulties in drawing generalized conclusions from our observations.

We believe that three key common phenomena of the intermediate zone indicate the feasibility of real-time monitoring of the treatment response: (1) antiphase structuring at the tumor visual and impedance boundaries; (2) local effects involving an activation and expansion of the IZ, and (3) systemic effects, characterized by the IZ subsequent reversion during the post-effect stage.

In summary, it seems appropriate to return to the definition of tumor boundaries by I. Prigogine as “an overlap in entropy production rate between the normal tissues and the tumor”. The discovery of DS around tumors may predetermine a paradigm shift in cancer research. From the perspective of DS theory, FBT can be conceptualized as thresholds beyond which the effects of weak stimuli are diminished or absent. By examining different scales—from whole tissue down to mitochondria—researchers can gain a comprehensive understanding of how DSs form and evolve.

Interpreting the presented phenomena through the lens of DS theory offers several advantages:

- It allows for a more comprehensive understanding of how complex systems behave under stress (like ischemia). Rather than isolating variables, it considers interactions and emergent properties;
- The DS theory emphasizes non-linear dynamics and how small changes can lead to significant effects, which is relevant when considering how tumors might respond to mild conditions;
- It provides insights into how biological systems can self-organize and adapt, which may explain the observed boundaries in impedance fluctuations as a result of cellular responses;

On the other hand, a biological-reduction approach may fall short because:

- It risks oversimplifying complex interactions within tissues by focusing solely on individual components without considering their interrelations;
- This approach often relies on static models that do not account for dynamic changes occurring during ischemia.

The presented findings demonstrate the potential of dynamic electrical impedance imaging to delineate tumor heterogeneity and offer novel insights into cancer-host interactions during targeted interventions in real time.

**Note:** *Some of the presented phenomena, such as antiphase effects and broadening of impedance boundaries during stimulation, can probably be partially confirmed using existing EIS time-lapse imaging tools and/or fluorescence microscopy samples. However, for a more adequate study of the SEL dynamics, as mentioned, a significant improvement in the spatio-temporal characteristics of the EIS technology is required*.

#### Additional remarks

The “order from chaos” concept is particularly relevant in tumor biology, where chaotic cellular environments can lead to the formation and progression of tumors. The tumor microenvironment may initially appear chaotic, manifesting as irregular cellular arrangements, heterogeneous cell populations, and varying extracellular matrix compositions and signaling pathways. The impedance fluctuation fields reflect the dynamic electrical properties of tissues that can change in response to different physiological states or stimuli. In this context, DS are systems that maintain their organization through the continuous exchange of energy, information, and matter with their surroundings, leading to complex behaviors observed in SEL dynamics. This process is particularly relevant when considering the melanoma zone characterized by impedance fluctuation fields, where initial chaos may precede the emergence of structured tumor formations.

In the absence of external influences, and due to the relatively stable genetic and phenotypic traits, a tumor may be defined as a quasi-stationary structure exhibiting characteristics akin to a static entity over extended periods (months to years). However, *it is crucial to note that DS can only be identified in dynamic states. Without external stimuli or exposure to perturbations, these structures remain undetectable as they do not exhibit significant changes in their static form*. Moreover, in the absence of external stimuli or disturbances, these structures may be obscured by background metabolic activity. Upon the application of external influences—such as measurement microcurrents—the tumor’s behavior may shift from a quasi-stationary state to a more dynamic one. The more specific stimuli - ischemia or EMF - serve as triggers that reveal underlying structural organization within a chaotic landscape. These stimuli may induce changes in cellular behavior and tissue architecture, allowing for the identification of tumor segments that exhibit heightened sensitivity (Figs. 9,12) or are approaching a bifurcation state, which is crucial for identifying functional boundaries and understanding tumor dynamics. (This phenomenon parallels how sick or weakened organisms respond more acutely to environmental factors due to compromised systems.) The application of external stimuli not only reveals but also contrasts various tumor segments based on their sensitivity to these influences. Tumor cells at morpho-functional boundaries, particularly at invasive fronts, are often more responsive to environmental changes due to their dynamic nature and adaptive capabilities. These boundaries represent critical zones where cancer cells transition from benign growth patterns to invasive behaviors, making them prime candidates for observation under weak stimulus conditions.

In case of the short stages of our experiments, the influence of the current metabolism on the IB “4” is seemingly insignificant* and therefore they can be considered as the boundaries of the effective zone of the current entropy/negentropy flow and - in the first approximation - as the minimal FBT. The presented images (Fig. 4,5,6,7,10,11,13) indicate that dimensions of the IZ significantly exceed the VB. The weak stimuli used can lead to a noticeable further expansion of IZ, and at different levels of cellular organization this expansion can differ significantly (Figs. 4,5,7,10,11).

* This is confirmed by the high reproducibility of the SEL in the absence of influences.

The observation of autowave processes and antiphase structuring around the tumor boundaries (Fig. 4) serves as compelling evidence for their dissipative origins, demonstrating that biological systems can spontaneously organize into complex patterns when subjected to weak stimuli. From the point of view of cellular biology, this effect can be explained by the tumor acidic microenvironment (Warburg effect) which is known to influence various cellular processes, including cell signaling, proliferation, and apoptosis. Cancer cells possess distinct bioelectrical properties characterized by a more depolarized membrane potential compared to their healthy counterparts^81^. The depolarization and hyperpolarization events occurring in cell membranes indicate a complex interplay between tumor cells and their microenvironment, which may lead to modifications in ion channel activity and signaling pathways within both intracellular and extracellular environments. This was also registered as a transit distortion of the ion balance not only around the visible border of the tumor, but also inside it (i.e. PN), Fig. 4h.

From the point of view of the DS theory, following a brief ischemic event, a negentropy flow is initiated in the most aggressive region of the tumor. This flow induces a wave of ordering characterized by synchronized changes in the electrical impedance of cell membranes within the peritumor zone. Such synchronization reflects an emergent property typical of DS, where local order arises from energy/information dissipation, allowing for enhanced cellular communication and coordination. The propagation speed of this wave may depend on several factors, including the intrinsic properties of the cell membranes (such as ion channel dynamics), intercellular connectivity, and the biochemical environment that influences signal transduction pathways.

The case shown in Fig. 6 demonstrates the existence of another mechanism for the emergence of an antiphase structure at the tumor border in the form of an expansion of the area of the original microclusters in response to ischemia. (Basically similar dynamics was observed in reponse to mm-EMF in another case^35, Fig.15^ shown in Fig. 1).

**Fig. 15.**
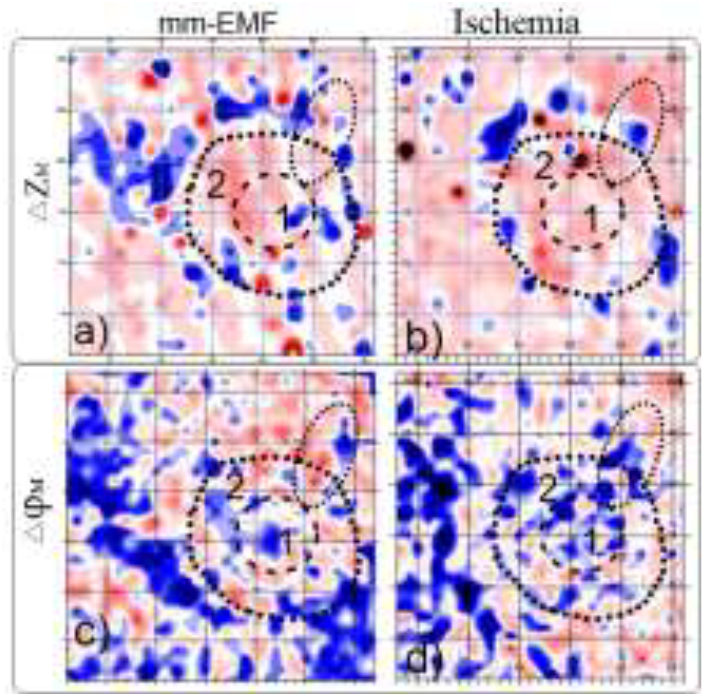
An example of the similarity between manifestations of weak information (EMF) and energy (ischemia) influence: **a**,**b)** Only on the outer contour of the tumor (cytosol level); **c**,**d)** On the outer and inner contour (mitochondrial level). Difference images of Z_M_ (up) and φ_M_ for the corresponding stages. *Both phenomena can be generalized under the concept of ‘order out of chaos’. They illustrate how biological systems maintain homeostasis or adaptively reorganize when subjected to stressors such as ischemia or non-thermal EMF. This suggests that both stressors elicit comparable biological responses across different cellular environments. Understanding these responses could have significant implications for developing treatment strategies targeting how tumors react to various forms of stress*.

The k-landscape configuration and its dynamics (Fig. 6) are presumably related to the metabolic plasticity of melanoma cells and their crosstalk with the surroundings. It alters ionic balance through lactic acid production, affecting ion transport, and is closely tied to mitochondrial dysfunctions impacting bioenergetics^76^. Thus, it may indicate spatial directions for oncogenesis through ionic imbalance area “5”.

In tumors like melanoma, cells often form microclusters or aggregates, enhancing survival under adverse conditions such as ischemia. These clusters can exhibit heterogeneous metabolic profiles, allowing some cells to survive even when others do not. Evidence supports that acidic regions surrounding melanoma are localized or clustered rather than randomly distributed throughout the tumor microenvironment. The presence of these clusters can affect the ionic balance in surrounding tissues due to altered secretion profiles (e.g., release of lactate, protons) and changes in membrane permeability. The post-ischemia expansion of clusters at the cytosol level shown in Fig. 6 further emphasizes the adaptive nature of cancer cells following stress events. Such expansions could lead to enhanced metabolic activity or altered signaling pathways that support tumor growth.

Magnetic resonance spectroscopy (MRS) visualized pH and exosome variations within tumors, revealing distinct clusters of acidity associated with aggressive tumor phenotypes^77^. Additionally, histological analyses have shown that these acidic zones often coincide with areas exhibiting high cellular density or necrosis, suggesting a relationship between metabolic activity and spatial organization within tumors^78^. These morphofunctional differences between the acidic regions surrounding the melanoma (Fig. 6c and Fig. 14b) and those further from the tumor edge may lead to contrasting reactions, such as the multidirectional response to hypoxia and an expansion of this zone in the impedance landscape in response to reoxygenation, which could potentially be used to identify them.

Understanding the localized nature of acidic areas surrounding melanoma and their sensitivity to short-term hypoxia has significant implications for treatment strategies. Targeting these specific microenvironments could enhance therapeutic efficacy by disrupting the adaptive mechanisms employed by cancer cells under stress conditions.

The initially expressed features of the quasi-static mitochondrial landscape (Fig. 7b) also manifested in its fluctuation fields (Fig. 7) in response to ischemia, as phenomena of advancing functional boundaries and increasing, then subsequently smoothing, the level of fluctuations. The sharp increase in mitochondrial impedance fluctuations during short ischemia reflects adaptive responses at both cellular and systemic levels within the peritumoral zone. From a cell biology standpoint, these changes can be related to local effects of mitochondrial fusion and fission under stress. In the DS context, the peritumor zone is characterized by increased fluctuations where energy is utilized for cellular communication and adaptation—characteristics of a system striving for stability amidst chaos. The circular configuration of mitochondrial impedance boundary “3” (Fig.7b) may be related to their tendency to migrate to areas of greatest energy demand. This area is typically more responsive to environmental changes due to its proximity to healthy tissue, which may still receive some blood supply. Also, neoplastic mitochondria may exhibit heightened sensitivity due to existing dysfunctions caused by mutations or altered signaling pathways associated with cancer metabolism. The propagation of fluctuation fronts signifies an important aspect of intercellular communication that may influence tumor behavior and recovery mechanisms following ischemic events. Enhanced mitochondrial fluctuations may facilitate communication between tumor cells and their microenvironment, potentially influencing tumor growth and metastasis. Conversely, the smoothing effect within the tumor core “1”and zone “4” indicates that tumor cells may have reached a threshold where further adaptation is limited due to extreme hypoxic conditions or metabolic shutdown. Despite no change in amplitude, this smoothing can imply metabolic suppression and loss of heterogeneity. (It would be very interesting to confirm this, only now discovered, functional similarity between “1” and “4” using appropriate laboratory methods.)

The process of ordering/disordering of the φ_M_ landscape is also manifested in the correlation fields of Fig. 8. It is evident that the maximum synchronization and the corresponding contrast of the φ_M_ landscape occur at the post-reoxygenation stage with subsequent rapid desynchronization. The significance lies in understanding how tumors adapt to microenvironmental stresses, such as hypoxia.

We were particularly impressed by how purely informational influences, such as non-thermal mm-EMF and weak MF, elicited profound changes in the impedance landscape. These responses challenge traditional views on electromagnetic interactions and underscore a novel understanding of information transfer in biological systems.

Against the background of the mitochondrial landscape (Fig. 7b), the “2+” zone stands out for its similarity to the melanoma nucleus “1”. The end-to-end graphical (Fig. 9) and subsequent fluctuation analysis (Fig. 10) revealed that this similarity is due to the high sensitivity of these areas to most of the effects used. In response to mm-EMF, this similarity manifested itself even more contrastingly in the φ_M_ fluctuation fields (Fig. 10) - in the form of two related structures: the PN stroma and the “daughter” structure “2+”. These areas can exhibit a heightened response due to the presence of reactive stromal cells and altered extracellular matrix components that influence ion channel activity and cellular excitability^79^. Moreover, the identified phenomena may be associated specifically with mitochondria as mobile, dynamic, energy-transforming organelles that transmit signals to all cellular compartments and the entire organism^80^. *This may have profound implications for understanding tumor biology and treatment approaches. The enhanced spatial dynamics suggest that these regions may serve as critical zones for therapeutic interventions, potentially allowing targeted approaches that exploit their unique electrophysiological properties. Furthermore, the ability of these regions to rapidly turn on and off in response to EMF suggests dynamic adaptability that may influence tumor behavior, growth patterns, and responses to therapy. In terms of DS theory, the results imply that FBT are not simply passive but actively engage in information exchange with external stimuli such as EMF. In our context, the enhanced spatial dynamics associated with EMF exposure may reflect a basic process of self-organization in these sensitive regions, leading to emergent properties that may influence cellular communication, metabolic processes, and overall tumor dynamics*.

Another striking manifestation of mitochondria as the DS phenomena is the existence of initial φ_M_ fluctuations (Fig.9b): according to the DS basics^36,54^, initial irregular oscillations that evolve into substantial and stable alterations in a system’s behavior due to minor disturbances are indicative of the system’s dissipative characteristics. This phenomenon highlights how small influences can lead to significant changes in dynamic systems (Fig.10). In the context of post-ischemic conditions, the cellular and mitochondrial responses within the stroma of a PN can be understood through several biological principles. Ischemia affects cellular metabolism and mitochondrial function. The observed gradual weakening of the nevus core’s landscape may reflect alterations in mitochondrial impedance, indicative of changes in bioenergetics and cellular homeostasis. Mitochondria play a crucial role in energy production and are sensitive to environmental changes; thus, their impaired function can lead to differential relaxation rates between internal (neoplastic) and stromal mitochondria. This disparity could suggest that the neoplastic cells are undergoing stress responses due to ischemic conditions, while stromal cells might be responding differently based on their functional roles within the tissue microenvironment.

From the perspective of cell biology and DS theory, these observations can be interpreted as manifestations of complex adaptive systems where biological entities maintain order through energy dissipation. In this case, the post-ischemic response may represent a shift towards a new equilibrium state for both neoplastic and stromal cells as they adapt to altered metabolic demands and oxygen availability. The differences in mitochondrial behavior could indicate varying capacities for adaptation among cell types within the tumor microenvironment, which is critical for understanding tumor resilience and potential therapeutic targets.

In response to the mm-EMF/MF switching on and off (Fig. 11), two different scenarios of the correlation fields restructuring emerged at the mitochondrial and cellular levels. Interpreting these scenarios solely through the lens of cell biology is difficult; however, they can be conceptualized as examples of “order from chaos”:

- Emergence and reversal of the mitochondrial front as local coherent - positive and negative - spatial correlations. This demonstrates establishment of localized order amidst global disorder;
- Dual ordering of the correlation field in response to EMF and MF, characterized by a predominance of high positive correlation in the background, juxtaposed with isolated regions/clusters of negative correlation at the peritumor area, Fig.11e-h.
- Switching off EMF revealed the opposite effects - a decrease in the background coherence (Fig.11g) and the *phenomenon of reverse migration of the negative clusters to the tumor border* (Fig.11i-k);
- The following MF exposure resulted in return of the high correlation of the background plus an additional peritumor zone that was “freed up” by the displacement of negative clusters.

From the perspective of the DS theory, the chaotic fluctuations in the peritumor zone give rise to islands of negative polarity, while the melanoma zone exhibits a transition towards higher positive correlation. This phenomenon exemplifies “order from chaos,” as localized interactions and energy/information exchanges within the system lead to emergent patterns and structures that enhance biological organization amidst an otherwise disordered environment. Each scenario reflects different aspects of how dissipative structuring operate under varying conditions of order and chaos while suggesting mechanisms through which negentropy flows might manifest differently depending on structural organization at multiple biological levels. The emergence of high positive correlation in one region indicates a robust ordering process, likely driven by external energy inputs or internal feedback mechanisms that stabilize certain configurations. Conversely, the presence of islands of negative polarity in the peritumor area suggests localized disruptions or *competing dynamics within the overall system*. These negative correlations may arise due to factors such as resource depletion, competitive interactions among cellular populations, or differential responses to environmental stimuli. This duality reflects the non-linear nature of dissipative structures where regions can coexist with varying degrees of order and disorder. In summary, the interplay between ordered and disordered states in this context illustrates how complex systems can evolve dynamically under specific conditions, leading to heterogeneous patterns that are characteristic of dissipative structures.

*There are analogs in biological systems where similar dynamics occur, e.g*., *(i) in neural tissues, individual neurons exhibit fluctuating electrical activities while simultaneously contributing to coherent patterns across networks; (ii) cardiac myocytes demonstrate synchronized contractions driven by localized electrical impulses but also maintain overall coherence necessary for effective heart function. These examples illustrate how both fluctuation-based dynamics (like those seen in mitochondrial ensembles) and coherence-based dynamics (as observed in intercellular communications) coexist in complex biological systems. In the context of the DS approach, while intercellular contents contribute to negentropy flow, their mechanisms differ from those of mitochondrial ensembles due to their reliance on extracellular interactions rather than intrinsic metabolic processes*.

Comparing the φ_M_ and Z_k_ patterns (Fig.11) suggests that although spatial electrophysiological responses are expressed differently—mitochondrial responses are more pronounced in fluctuation fields due to their localized nature, while intercellular content responses are more pronounced in coherence fields due to collective behavior among cells—the underlying principles governing these processes remain interconnected through their contribution to overall cellular organization.

The emergence of structures around the tumor and within the stroma under EMF exposure indicates a form of DS, where non-equilibrium conditions lead to self-organization and increased spatial fluctuations in impedance. This phenomenon suggests that mitochondria can adaptively respond to environmental stimuli by reorganizing their functional architecture, thereby enhancing their capacity for energy production and cellular communication. Moreover, the transient nature of these structures upon cessation of EMF exposure implies a dynamic equilibrium between stability and fluctuation inherent in biological systems. The ability of these structures to reappear with renewed EMF exposure while maintaining their shape signifies an underlying resilience characteristic of dissipative systems. In contrast, the cytosol’s rapid but singular response may reflect its role as a more immediate mediator of cellular processes without the complex feedback loops present in mitochondrial function. This disparity underscores the importance of mitochondrial dynamics in understanding how cells interact with external stimuli and adapt over time.

The analogy between ischemia (an energy-related effect) and mm-EMF (an information-related effect) lies in their respective impacts on mitochondrial function. While ischemia primarily disrupts energy supply and induces stress responses, the mm-EMF exposure appears to modulate cellular communication and information processing within mitochondria. Both stimuli ultimately affect mitochondrial dynamics but do so through different mechanisms—ischemia through energy depletion and mm-EMF through informational modulation—highlighting the intricate interplay between bioenergetics and cellular signaling.

The phenomenon of reverse migration of the negative *clusters* (Fig. 11i-k) could be explained through cell biology’s insights on cellular response mechanisms combined with the principle that DS can undergo phase transitions based on external conditions^36,48,52,53^. When an external influence is applied, it may drive the system into a new state (e.g., tumor growth), but once that influence is removed, the system may revert back to its original state or another stable configuration. The reversible dynamics observed across various natural phenomena illustrate fundamental principles governing both living organisms and physical systems:

- *Cells exposed to growth factors proliferate; upon removal of these factors, they may cease division)*.
- *Neurons exhibit synchronized firing patterns under certain stimuli but return to baseline activity when those stimuli are removed)*.
- *Under nutrient-rich conditions, individual amoebae aggregate into multicellular structures that exhibit coordinated movement and function—a dissipative structure formed through collective behavior. When nutrients are depleted (removal of external influence), these structures disband back into individual cells)*.
- *Ecosystems, like e.g. coral reefs, often display dissipative characteristics where species interactions create stable communities Changes in environmental factors can lead to shifts in community structure that may revert if conditions stabilize again*.
- *The Belousov-Zhabotinsky reaction showcases how chemical systems can form spatially organized patterns (DSes) under certain conditions but revert back to homogeneity when reactants are depleted or conditions change)*.

The weak ordering effects revealed in the X-ray irradiated melanoma (Fig.12-14) seem linked to polarization and depolarization at membrane interfaces, which may play a crucial role in maintaining some level of coherence despite the overall disruption caused by radiation. Radiation therapy appears to stabilize the fluctuation fields of impedance within cellular fluids and mitochondria, suggesting a modulation of cellular bioenergetics. Despite this stabilization, cellular membranes maintain their integrity, as evidenced by their continued responsiveness to mm-EMF and MF. This indicates that while radiation therapy induces significant changes at the cellular level, it does not compromise the fundamental properties of cell membranes, allowing for ongoing interactions with external electromagnetic influences. Cell membranes might show less change relative to others since they serve primarily a protective role but still experience indirect effects through oxidative stress from free radicals generated during radiolysis. While polarization/depolarization alone may not be sufficient for coherence without proper coupling mechanisms (like gap junctions), it can contribute significantly when combined with other factors^82,83^.

The tumor microenvironment can be conceptualized as a system of coupled oscillators where cancer cells interact with surrounding stromal cells, immune cells, and extracellular matrix components. This model helps explain reciprocal interactions between tumors and their environments. In the melanoma impedance landscape, the initial fluctuation field represents a state where there are varying levels of electrical impedance across different regions of the tumor. The presence of the tumor as a cluster with maximum fluctuations indicates areas of heightened activity or instability within its environment. This cluster’s characteristics are crucial for understanding how it interacts with external influences.

When exposed to weak influences such as non-thermal EMF and MF, the dynamics of the entire correlation landscape undergoes a transformation towards uniform coherence. This shift implies that even minor perturbations can lead to widespread changes in system behavior, aligning with principles of self-organization where small inputs can yield large-scale effects.

*In living organisms, similar discrete transitions can be observed:*

- *Neural Synchronization (In neural networks within the brain, neurons often transition between chaotic firing patterns and synchronized bursts depending on external stimuli or internal states)*.
- *Population Dynamics (In ecology, species populations may undergo sudden shifts (catastrophic shifts) due to environmental changes—akin to bifurcations seen in physical systems)*.
- *Circadian Rhythms (Organisms often display discrete shifts between different activity levels based on light exposure or other environmental cues)*.

Regarding tumor entropy emitters—hypothetical constructs referring perhaps metaphorically to tumor behavior under treatment—radiation therapy’s impact could theoretically lead either toward an increase or decrease in “emitter power” based on how effectively it disrupts tumor growth dynamics versus inducing adaptive resistance mechanisms within tumors.

However, empirical evidence specifically addressing this concept remains sparse; thus definitive conclusions cannot be drawn without further research into this area. Unfortunately, we do not have data on the state and size of the IB before irradiation, what could shed light on this issue. By measuring variations in impedance landscape before and after irradiation, researchers can detect shifts in entropy flux that result from changes in cellular activity or microenvironmental conditions.

In summary, the influence of X-rays on reciprocal cancer-host interactions is profound, leading to significant alterations in cellular dynamics characterized by the weakening or disappearance of autowave and coherent structures within melanoma zones. Understanding these changes provides valuable insights into potential therapeutic strategies as well as methods for visualizing and monitoring these complex interactions.

The presented series of spatiotemporal structuring phenomena correspond to such basic concepts of dissipative structures as non-equilibrium conditions, self-organization, bifurcation points, entropy/information production, interdependence with external factors. This analysis not only elucidates the intricate dynamics underlying the SEL but also highlights the critical role these phenomena play in understanding tumor behavior. The observed similarities in antiphase structuring across different biological entities—malignant melanoma, benign nevi, and plant leaves—indicate that such patterns may transcend species boundaries, hinting at fundamental principles governing biological responses to external stimuli. These findings underscore the importance of considering both physical and informational influences when assessing tumor characteristics and their functional boundaries. This insight opens avenues for further research into the underlying mechanisms that could unify our understanding of biological systems across various forms of life.

It is important to understand that in real time (seconds to minutes) DS in the tumor area can be reliably identified only in states of altered dynamics, since they do not demonstrate significant changes in their static form. Moreover, in the absence of external stimuli or disturbances, these structures may be obscured by background metabolic activity. This is because the oncogenetic dynamics associated with entropy and negentropy flow occur over significantly different time scales, typically spanning months to years. The near field of the flow is apparently reflected in the SEL snapshots as quasi-stationary IB: (i) they are significantly wider than the VB; (ii) the test-induced changes occur precisely there. In case of SSM, significant spatial differences are noticeable in the zones of influence of emitters at the intra-/extracellular levels from that at the mitochondrial level. Thus, these quasi-stationary IB can be defined as morphofunctional, in contrast to the mobile, test-induced FBT (2+, 3+, 4+). The term FBT also includes punctate microscopic clusters (Fig. 5) that only become apparent in response to ischemia. The internal border (of the primary nevus) should probably be classified as a morpho-functional or morphological one (like the visible borders of the tumor), since it can be distinguished on the snapshot (Fig. 3). However, the boundaries of the PN (the stroma, Fig. 10), which manifested as a fluctuating structure only in response to the information impact (EMF/MP), should definitely be defined as a fluctuating internal boundary. Notably, the high-amplitude antiphase activity can also serve as a marker of the morphological and morphofunctional border since it can occur on opposite sides of this border or even twice in the metastasis zone (such as the visible and hidden satellite metastasis of MS, and zone “2+” of SSM). The influence of these most powerful emitters of entropy/negentropy determines the configuration/asymmetry of the external morpho-functional and, accordingly, FBT.

#### Shortcomings and potential developments

The sample size is limited, with only five main experiments analyzed. In the context of the general acceptance and universality of self-organization and dissipative structure theories, this seems sufficient to represent the discovery of new phenomena. However, from a clinical perspective, given the complex biology of melanoma and the significant individual differences among patients, a larger data set is needed to justify the generalizability of these results. This would not only increase the statistical power of the study, but also provide a more complete understanding of how these tissue-level dissipative structures manifest in different tumor cases.

Another limitation regarding the direct applicability of the observed phenomena to oncology is the absence of direct biological validation for the proposed dissipative structures. While this study reveals compelling electrical and functional dynamics within melanoma tissue, it does not provide definitive molecular or histological correlations with established mechanisms of tumor biology.

Additionally, while the results suggest that weak EMF and MF exposures induce significant biological effects at the tissue level, it does not fully address potential confounding factors. The interactions between tumors and EMF/MF remain a contentious area in medical science, and the further study would benefit from more rigorous control experiments to rule out alternative explanations for the observed impedance changes.

The current methodology is constrained to the examination of surface layers of tissue. To enhance its applicability, it is essential to incorporate a tomographic mode that would allow for deeper imaging capabilities. Furthermore, improvements in sensitivity and speed are necessary to obtain more accurate and timely results. Investigating the electrical impedance dynamics of deeper tissue structures necessitates the adaptation of alternative imaging techniques, with MRI being a primary candidate

To advance these studies, employing both 2D and 3D cell models will be crucial. Additionally, utilizing fresh tissue samples alongside dynamic visualization tools such as fluorescence imaging and MRI will likely yield the most promising results for understanding the complexities of melanoma at a tissue level.

Collaborations between oncologists, physicists and biophysicists, and medical imaging specialists will be vital for translating the presented pioneering discoveries and theoretical concept into practical diagnostic tools.

## Abbreviations

EIS: Electrical impedance spectroscopy
(EMF): Electromagnetic
(MF): magnetic fields
ECF, ICF: Extra- and intracellular fluids
DS: Dissipative structures
FBT: Functional boundary of a tumor
IZ: Intermediate zone
IB: Impedance boundaries
IB: Ion imbalance index
SEI: Skin Electrodynamic Introscopy”
SEL: Skin electroimpedance landscape
SEL: Spectral impedance landscape
TME: Tumor microenvironment
VB: Visible boundaries

## Data availability

The dataset for this article can be found online at DOI 10.6084/m9.figshare.28540361. It includes Supplementary Material and raw data of time-lapse image matrices in .txt format

## Acknowledgements

Sincere thanks to Prof. T. Dandekar, Dr. G. Dandekar, Dr. A.Riedel and Dr. Florian K. Groeber-Becker for their understanding and assistance, without which this work in wartime conditions in our country would hardly have been possible. Special thanks to Prof. V. Maksymenko and European Association for Cancer Research (Ukr-Award 04, 2022)

## Supplementary materials I

**Figure S1.**
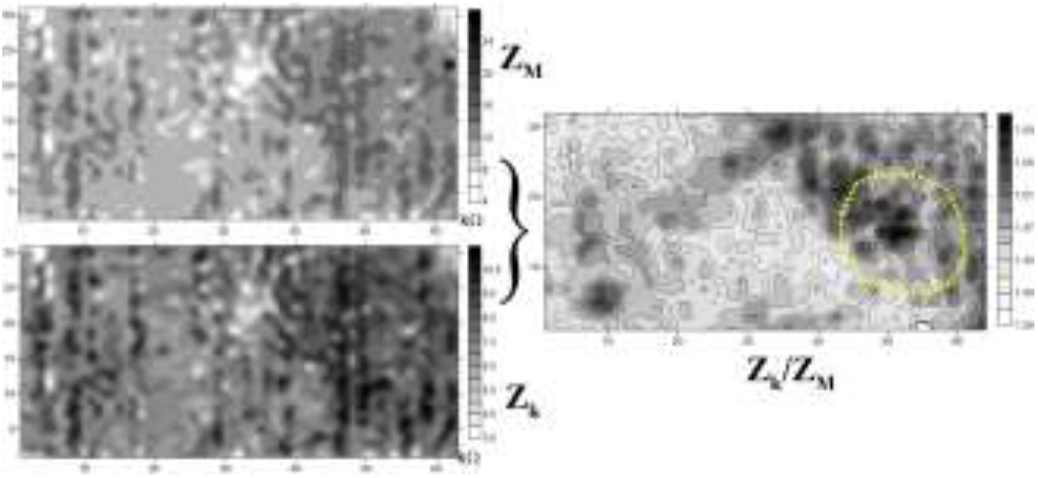
Raw images of the intra- and intercellular impedance module, which show the structural distortions of the primary transducer, and their ratio CDI =Z_k_/Z_M._.

**Figure S2.**
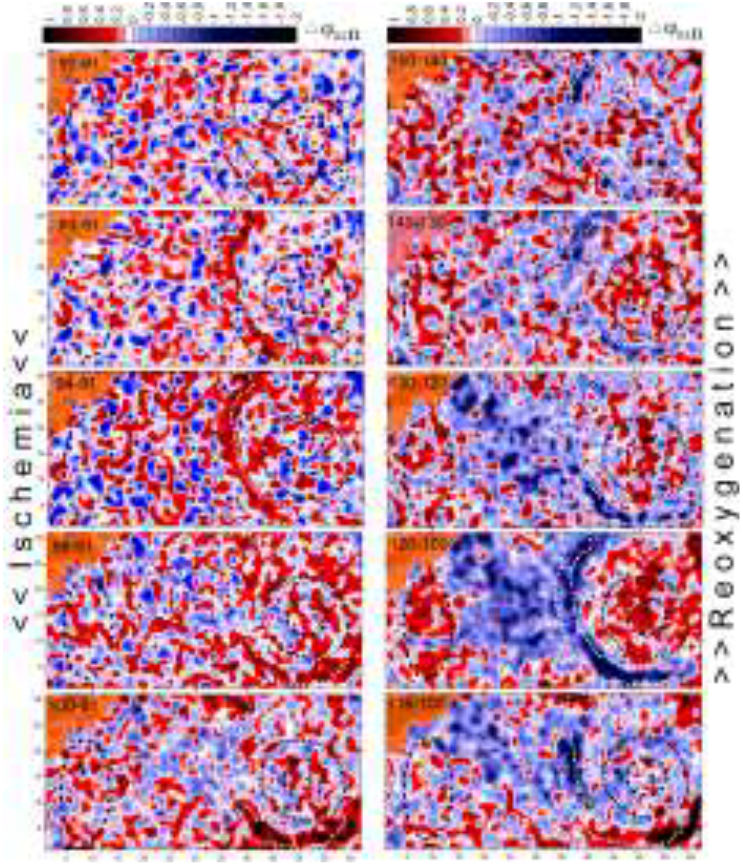
Temporal dynamics of the φ_M_ landscape during the ischemia-reoxygenation period as difference images. The “3+” structure as a circle begins to form already in the 3rd frame of the ischemia stage. The DS in the center of the frame is clearly emerges and disappears in the reoxygenation stage.

**Figure S3.**
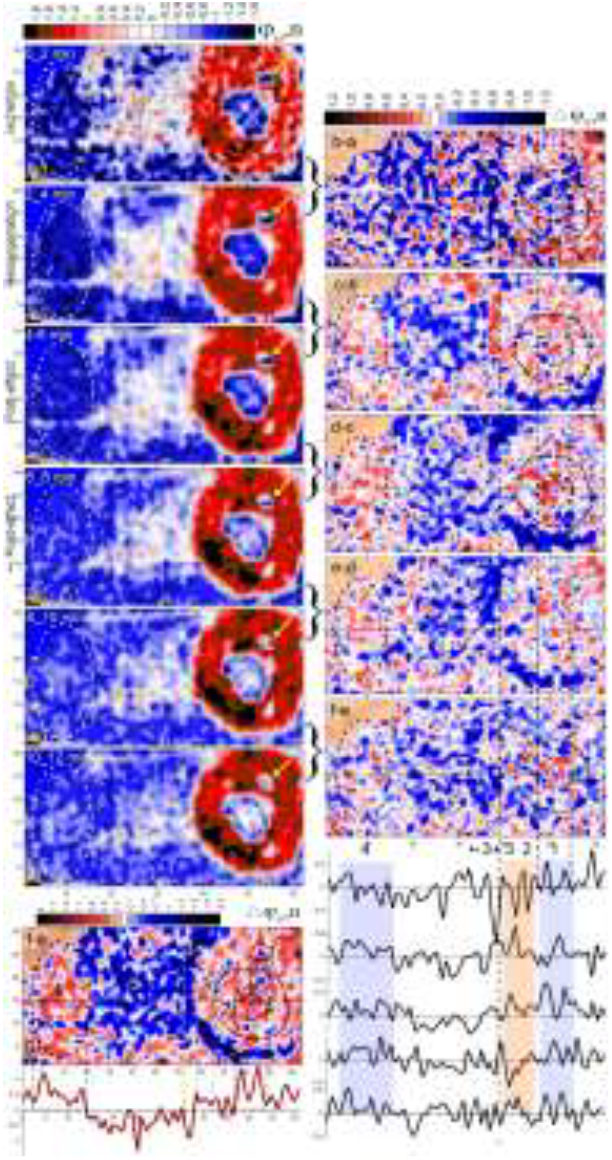
Step-by-step and endpoint changes of φ_M_ initial landscape during the entire ischemic test. **a**,**f)** Normalized φ_M_-maps averaged over 2-minute periods; **g)** Map of endpoint change. Right: differences of **a-b**,**b-c**,**…**,**f-e**. Bottom: the landscape profiles. Of note: Contrasting stroma of the PN during **c** and **f**.

**Figure s4.**
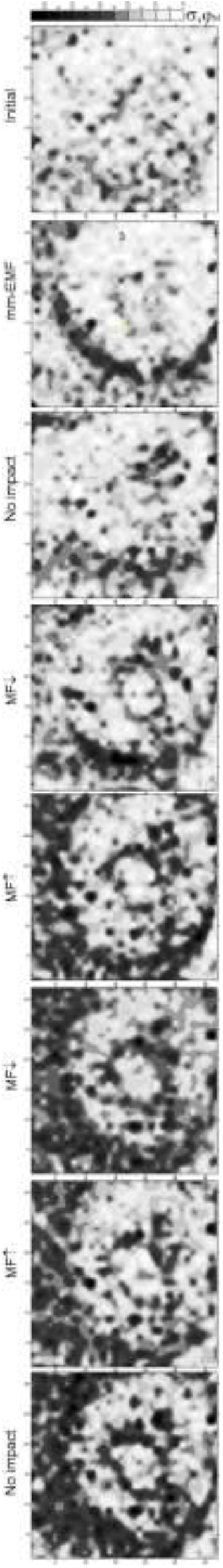
The φ_M_ spatial fluctuations reveal internal and external boundaries of SSM as the DS structures in response to mm-EMF and MF exposure

**Figure S5.**
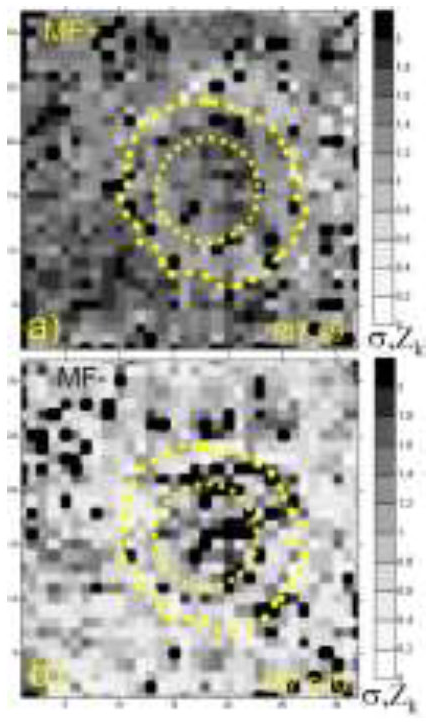
The phenomenon of weakening of fluctuations at the extracellular liquid level (Z_k_) in response to the MF turn-over. Dispersion fields (raw maps): **a)** MF↑, b) MF↓.

## Supplementary materials II

### <u>Analysis III</u>. Papillomatous nevus

This case was previously presented^33^ without analysis – just as an illustration of the antiphase structuring in the suspicious nevus. Here we present this case as a further evidence that the collective effects of ‘guest-host’ interaction, observed above in malignancy, may also manifest, to varying degrees, in cases of benign neoplasms. At the same time, due to the paucity of such observations, the analysis presented below should be considered a mechanistic study.

*The concept of functional tumor boundaries, while primarily associated with malignant neoplasms, can be considered in the context of benign nevi. These boundaries are not as clearly defined as in malignant tumors, but there are mechanisms that limit the growth and progression of nevi, effectively creating a functional boundary*^*89,90*^.

*In contrast to melanomas, which exhibit increased cellular disorganization and chaotic growth patterns, benign nevi display a more organized structure relative to surrounding tissues*^*68*^; *thus, they can be considered as emitters of negentropy. In this case, papillomatous projections can be considered more powerful entropy emitters due to their altered metabolic states and increased activity levels associated with viral replication. Such cells often exhibit heightened sensitivity to external stressors like EMF and ischemia*^*69*^ *due to their compromised homeostasis*.

*Therefore, while the concept of FTB in benign neoplasms like papillomatous nevi differs from that in malignant tumors, it is still relevant in defining the lesion, understanding its interaction with surrounding tissues, and guiding clinical management*.

In contrast to the melanoma studies, among 17 nevi examined, there were only 2 cases in which the impedance boundaries (IB) significantly protruded beyond the visible boundaries (VB) of the lesions.

In the case under consideration, the unusualness of the IB manifested itself in the initial SEL not only in their noticeable excess over the VB, but also in the form of two pairs of finger-like branches of the SEL extending from the IB and significantly exceeding it in amplitude (Fig. S12A, B). The most intriguing fact was that the dynamics of these structures was synchronous with the adjacent segment of the IB (Fig. 1, Fig. S12a), which further confirms their related origin.

Within the framework of the hypothesis of invasive/active fronts as entropy emitters, this ensemble can be represented as a 3-component emitter with characteristic antiphase dynamics of the outer sides. At the same time, depending on the degree of activation of the emitter, its area will also change. However, unlike locally located emitters in the above SM and MSS, due to the focusing arrangement of this one, the resulting flow of entropy will be concentrated mainly between them, thus creating a specific asymmetry of the outer SEL.

The experimental protocol was similar to the above cases, except that just a breath-hold of ∼40 sec was used as ischemic test, Fig. S13.

### 3.1. Information vs force impact: similarities and differences in coherent structuring

*When penetrating host cells, HPV viruses cause changes primarily in cell membranes, which is due to their effect on the activity of ion channels and cellular signaling mechanisms*^*73*^.

Fig. 12B shows the initial snap shot landscape in the parameters of cell membranes, φ_k._ Dark areas (marked as “2” and “4”), spreading from the visible border of nevus “1” may represent HPV-infected areas^70^, i.e. the supposed main emitters of negentropy. To demonstrate similarity of the response of φ_k_ to the applied stimuli, its dynamics presented as two corresponding columns of correlation fields, Fig.12a,g.

Maps (**a-e**) represent initial stages. The most significant initial dynamics are presented as a complex of (blue) structures “2, 1+,4” surrounding the (red) antiphase zone “3”. Exactly the same configuration was restored after the mm-EMF test in (e): only a slight increase in the contrast between the red and blue structures of the complex can be noted after the whole mm-EMF+MF test (stages 2-5, Fig.13). It is also important to emphasize that only a part of the nevus contour stands out in the most contrast, i.e. “1+”.

General similarity. In both columns (**b**,**f**) and (**c**,**g**), one can see a biphasic response to the stimuli, which, to simplify, can be expressed as “red rising absorbs blue (except “3”)” and vice versa: **b**,**f**) The correlation coefficient ***r*** across the entire impedance landscapes increased reaching a value of 1 in **f**, indicating a high degree of order. However, in the projections of the nevus, this coefficient was negative (−0.66 *≤* ***r*** and -0.38 *≤* ***r***, respectively), suggesting localized dysfunction. In this case, the ordering of the surrounding landscape “**f”** occurred with **r** ≈1.0 almost everywhere, whereas in **“b” –** only in “3”, i.e. in the area of the emitters focusing area.

**Fig. S12.**
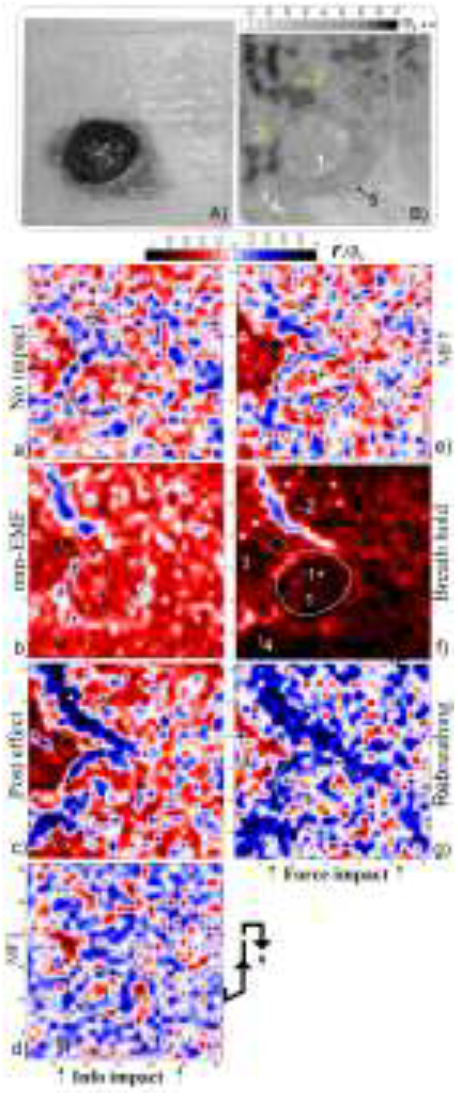
Information vs force impact on the collective dynamics of cell membranes. **A**,**B)** Photo of the nevus and a snap-shot of the φ_k-_landscape; Correlation fields of φ_k-_ landscapes: **a**,**e)** Initial stages; **b**,**f)** The mm-EMF exposure and ischemia; **c-g)** Post-effects; **d)** The MF exposure (p<0.01).

**Fig. S13.**
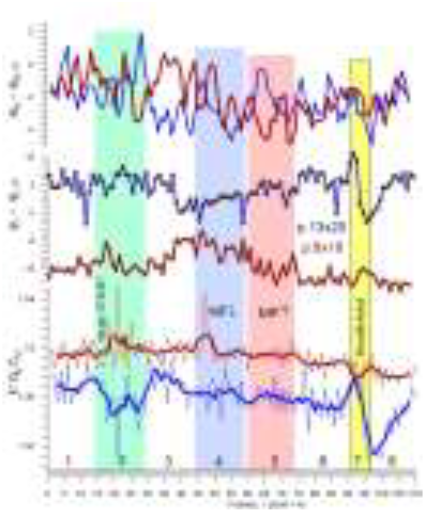
Counter-directional dynamics throughout the experiment: time graphs of pixels 8×18 and 13×20. Bottom to top: Ionic balance k; deviations of φ_k_ and φ_M._ Of particular note is the phenomenon of locally increased sensitivity of mitochondrial membranes (than those of cell membranes) to EMF/MF exposure, which was also registered in the SSM, Fig. 9. *Notably:* - Antiphaseness of *k* and φ_k_ in response to mm-EMF and ischemia; - Antiphaseness and sensitivity of φ_M_ everywhere only before ischemia with subsequent synchronization only before ischemia followed by φ_M_ syncronization.

**c-g)** The similarity of both patterns can be defined primarily as the ordering and expansion of the negative coherence zone (up to -1.0 ≤ ***r***) with the destruction of order outside the “2.1+4” sector, which is especially pronounced in the response to ischemia.

**(d)** The MF-induced destruction of order followed by its recovery in “e”.

### 3.2. Ischemia-induced dynamics and clusterisation

**Fig. 14 presents the time-lapse images of emergence, propagation and relaxation of the complex of coherent in-phase and out-of-phase structures at the level of cell membranes during the breath holding and subsequent restoration. The normalized difference images of the** φ_k_ landscape **highlight emergence and disappearance of “1+”, “2”, “3” and “4” structures approximately 8–12 seconds after breathing stops, Fig. 14c. The dynamics of all these structures appear to be synchronous: the blue structures “1+”, “2”, and “4” appear as a single in-phase ensemble; the red out-of-phase structure “3” located between them, and, only much less pronounced redness, everywhere (including the most of the nevus) on the opposite side of the paired emitters “2” and “4”**.

**The structures “1+” and “3”, as in the response to mm-EMF, are located on different sides of the visible border of the nevus thereby opposing *only* its active (viral) part “1+”**.

**Comparing Fig. 14c,d, one you can assess the initial velocity of the red and blue structures of the entire “2-4” complex as 1-2 mm/sec, i.e. similar to that of MS (Fig.2)**. This value exceeds **that observed for calcium wave propagation since the dynamics of ion movement leading to collective membrane depolarization occurs at a higher rate**. However, already in the next frame it can be noted that this speed decreased approximately by half (Fig. 14d). In addition, it is noticeable that on the outer side of the emitters “2,1+,4” the speed of propagation of blue structures is noticeably lower.

The delays and abnormalities in intercellular communication during these phases can alter ionic balance within tissues. As signaling cascades unfold over time, variations in ion concentrations (such as Na^+^, K^+^, Ca^2^+) across membranes may occur due to differential permeability changes and active transport mechanisms. These alterations can affect conductivity ratios between interstitial fluid and cytoplasm, influencing overall tissue excitability and responsiveness.

**Fig. S14.**
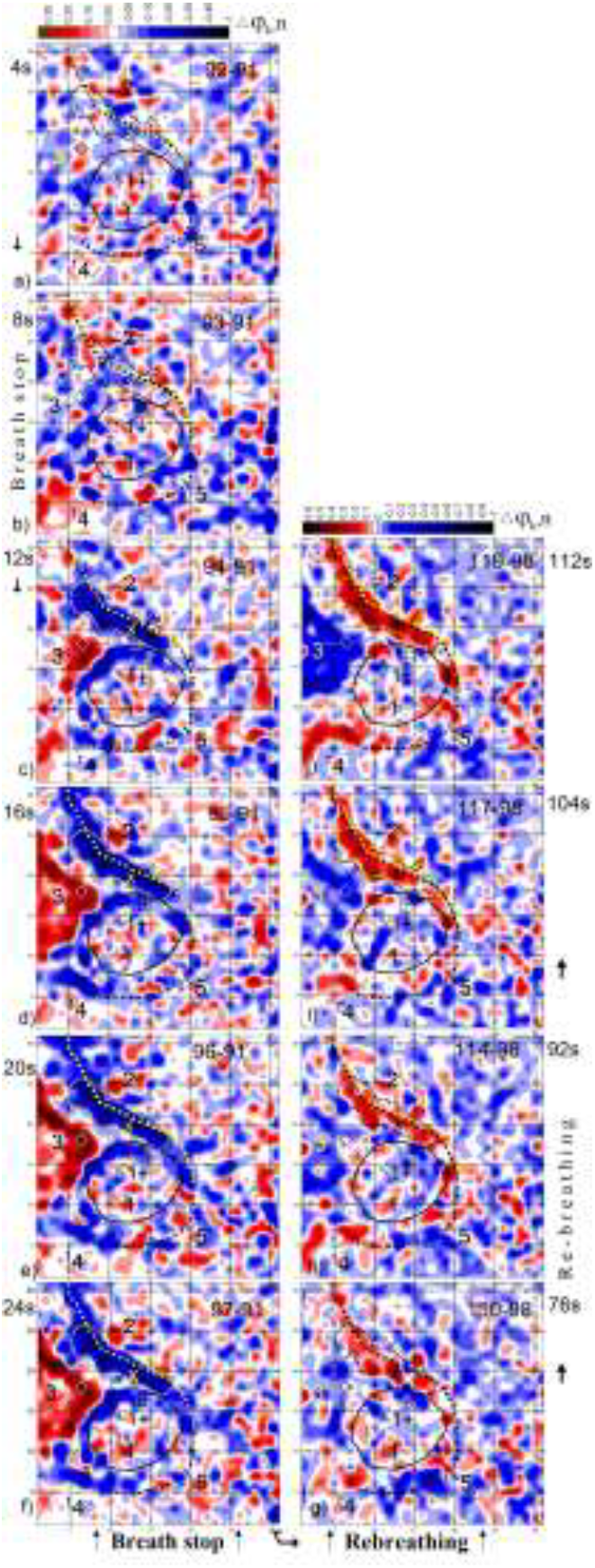
Initial dynamics of φ_k_ structures “1+”,”2”,”3” and “4”. Difference of normalised frames (93-91, 94-91, 95-91). □-p.13).□20;).□-p.8).□18

Fig. 15 demonstrates restructuring of the ionic balance landscape during the breath-holding period:

- Effect of ischemia on the distribution of the *k* index, where the blue structures can presumably correspond to the HPV infiltration map;
- Time dynamics of three characteristic points of the active zone “1+,2, 3”;
- Local changes in *k* in the “2-4” area can exceed the background values by an order of magnitude or more.

**Fig. S15.**
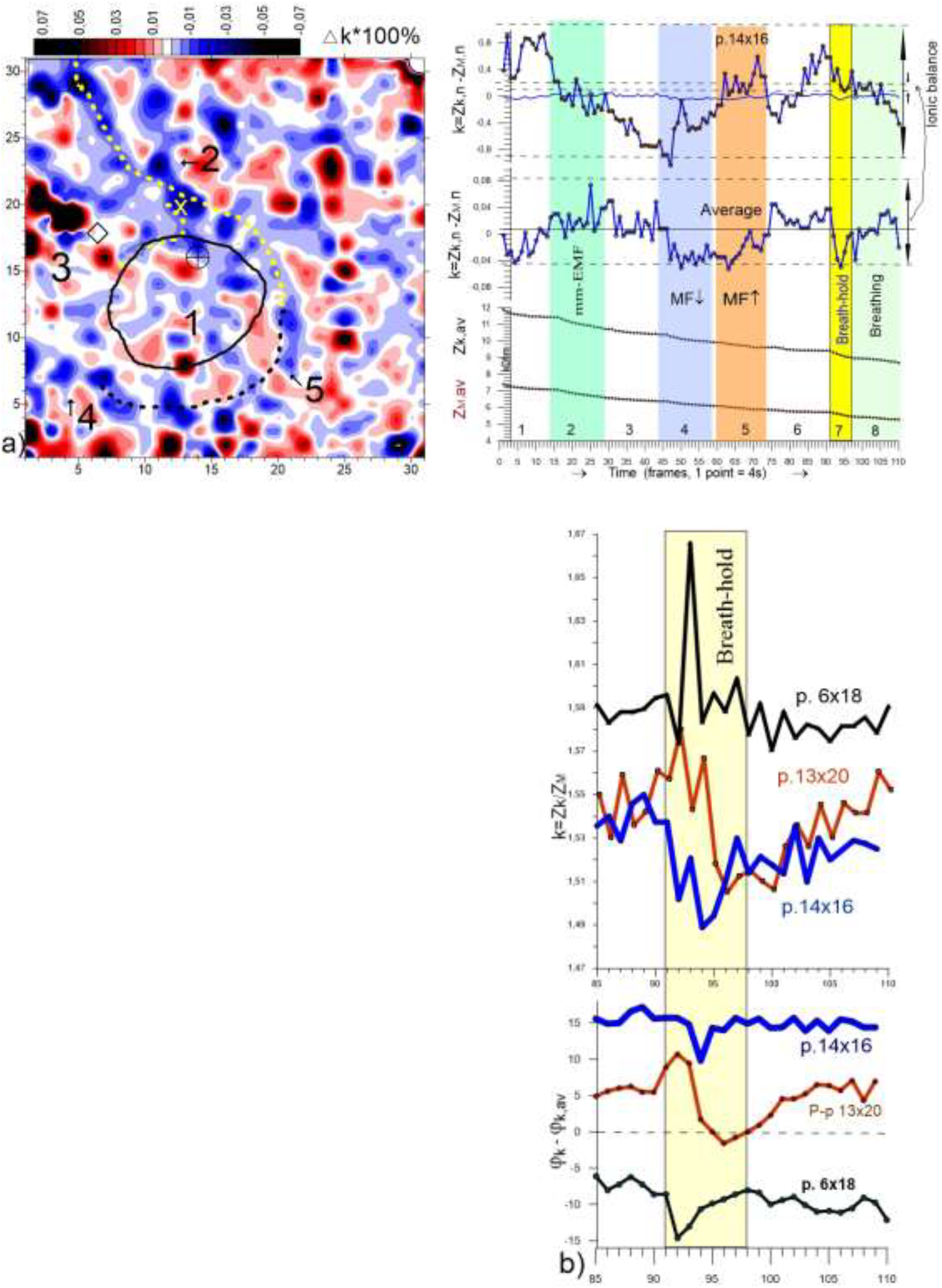
Restructuning of the ionic balance landscape during the breath-hold period. **Left)** Difference image of the ***k*** maps (99-91). **Right**) Temporal curves of ***k*** for the whole experiment: of a pixel 14×16, the frame-average fluctuations, and real current values of Z_k_ and Z_M_, p<0.001 □ - p.13□20; □-p.6□18;□-p.14□16 Note. (i) Increased sensitivity of the boundary segment “1+” (p. 14×16) to all stimuli; (ii) Higher sensitivity of the points of zone “2-3” (13×20, 6×18) to ischemia →.

***The main common feature with the above cases of melanoma is the expansion of FTB in the form of activation and spread of the frontal dynamics of the initial impedance landscape. In particular, it is worth noting the similarity in the following effects and phenomena:***

- ***Ischemia-induced antiphase structuring with propagating wave at the cell membrane level (Fig. 4);***
- ***Ischemia-induced effects of ordering/disordering of fluctuation field of the mitochondrial membrane impedance and advancement of the φ***_***M***_ ***functional boundary (Fig. 7);***
- ***Large diversity of temporal response of individual points (Fig. 9)* ;**
- ***Response to both: EMF and hypoxia (Fig. 9);***
- ***(less pronounced, but noticeable) Anomalies in the spatial distribution of ion balance (Fig. 6);***
- ***Increased sensitivity of boundary points to EMF and MF (Fig. 9);***
- ***Dual ordering of the correlation field in response to EMF and MF, characterized by predominance of high positive correlation in the background, juxtaposed with isolated regions/clusters of negative correlation (supposed DS) at the peritumor area, Fig. 11;***
- ***Restructuring of the ionic balance landscape during the breath-hold period, Fig.6f***.

The observed dynamics can be interpreted through several lenses: **oncogenic nature of HPV, quorum sensing or viral signalling effect, and** systemic responses following ischemic injury. The transition from positive to negative correlation coefficients can be indicative of malignancy within the nevus and its projections. While these observations do not serve as definitive evidence of malignancy on their own, they align with established principles regarding tumor biology and HPV’s role in oncogenesis.

Such dynamics can be also assessed as the initial compensatory mechanism of a two-stage scenario aimed at maintaining cellular integrity and function despite viral interference. In the subsequent phase, however, a pronounced antiphase reaction emerges within these projections, leading to a significant weakening of correlation in the surrounding tissue. The transition indicates a shift from compensatory responses to potential destabilization within the cellular environment. These findings suggest that virus-infected regions act as emitters of negentropy within a DS framework; they disrupt normal cellular function while simultaneously expanding under external stimuli. This duality highlights how pathological entities can exploit environmental pressures for growth despite their detrimental effects on overall tissue homeostasis. Moreover, during this second phase, characterized by the spread of antiphase structures, there is a disturbance of the surrounding landscape, which favors the increase of antiphase islands compared to EMF. This raises intriguing questions regarding underlying mechanisms: Firstly, it may be posited that an increase in the size of negentropy emitters correlates with diminished negentropy flows. Secondly, ischemia appears to exert a more profound influence on the HPV quorum sensing^71^ than EMF does, suggesting that cellular communication and collective behavior may be more significantly altered under ischemic conditions.

The rates and delays associated with these two phases are influenced by the number of cells in the ensemble. In larger cell populations, there tends to be enhanced intercellular communication through mechanisms like paracrine signaling or electrical coupling via gap junctions. This can lead to synchronized responses among neighboring cells, potentially speeding up both phases of the response.

Cell biologists study this dynamics using techniques such as e.g. fluorescence microscopy to assess changes in membrane potential or intracellular calcium levels during these phases^72^. DS theory helps explain how these coherent collective responses emerge from individual cell behaviors under external perturbations. The rapid primary response could be viewed as an emergent property arising from local interactions among cells. The slower secondary phase might reflect a transition towards new stable states as cells reorganize their internal structures and functions in response to prolonged stress.The delayed activation of certain pathways may lead to an extended area where coordinated responses occur as neighboring cells begin synchronizing their activities based on restored oxygen levels. This expansion reflects not only recovery but also potential alterations in tumor microenvironments that could influence tumor progression or treatment outcomes.

In conclusion, HPV is the primary cause of cervical cancer, a major global health concern affecting women worldwide. However, current detection methods often lack sensitivity, speed, or cost-effectiveness, especially in resource-limited settings^74^. Our findings elucidate a complex interplay between viral infection and cellular responses within peritumoral papillomatous nevi. The dual-phase reaction underscores critical shifts in cellular dynamics influenced by both ischemic conditions and negentropy considerations. Future research should further explore these mechanisms to enhance our understanding of tumor biology and potential therapeutic interventions.

### <u>Analysis IV</u>. Antiphase structuring and post-stress effect in a plant leaf*. Common features and differences from the above cases

*\* New results*

The scanning was performed on a succulent leaf (not separated from the plant) with a time resolution of 8 sec. A drop of laundry bleach was used as a stimulus. Fig. 16c,d shows the primary response to the stimulus in the parameters of the resistive and capacitive resistance of the extracellular environment, i.e. R=|Z_k_|cos(φ_k_), Xc=|Z_k_|sin(φ_k_). The difference images demonstrate the instantaneous manifestation of antiphase structures in both parameters around the site of stimulation. However, there are significant differences, in particular:

- spatial response in the form of a family of cell membrane mini-clusters (c) is also observed simultaneously in the vessel branching zone «2» due to the fast mechanisms of systemic signalling;
- the response of the extracellular environment (d) is more extensive, but there is an interesting nuance.The outline of this zone (indicated by the dotted line) coincides with the outline of the zone of reduced or zero response (c) (worth comparing with Fig. 3c1,2).

**Fig. S16.**
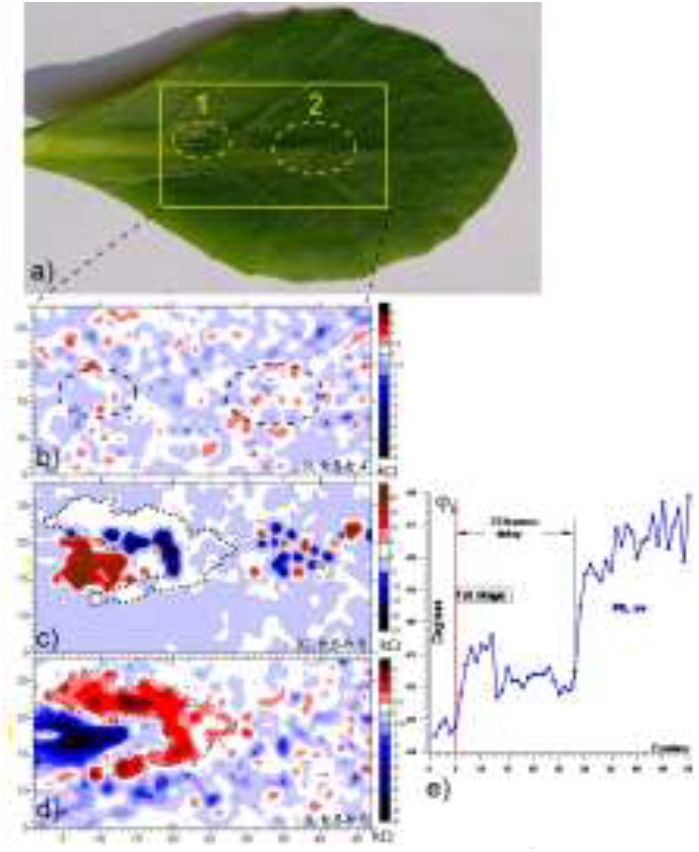
Restructuring of the impedance landscape in a plant leaf following localized burn and post-stress effect. **a)** Photo of a succulent leaf (still attached to the plant). “1” — burn area, “2” — vessel branching; **b)** Difference map of two frames of the previous frames of the resistive landscape, i.e.fr.5-fr.4 (as seen in e); **c**,**d)** The first appearance of the response of the extracellular capacitive and resistive landscapes, as fr.6-fr.5; **e)** Time plot of the average phase angle of the extracellular impedance, p<0.001.e

A description of all the results of this experiment is beyond the scope of this article, but it seems appropriate to note the phenomenon of a secondary stress response associated with the above-mentioned manifestation of the phenomenon of post-ischemic damage (Fig. 5). Fig. 16d shows time graph of the phase angle of the impedance of the extracellular environment, demonstrating this secondary, significantly more pronounced, systemic reaction that arose approximately 2 min after the end of the local reaction.

This situation can be also analyzed through the lens of DS^36,51^. In plants, when a leaf is irritated, it can trigger a spatial antiphase reaction, where areas of excitation and inhibition occur in a patterned manner around the site of irritation. The spatial antiphase reaction means that energy* is not merely consumed but transformed into organized patterns or structures. When an alkali drop irritates a leaf, it disrupts homeostasis and leads to localized changes in ion concentrations and membrane potentials. This disturbance can create gradients that spread across tissues to maintain order through coordinated reactions in different regions.

*\*The energy from heat causes denaturation of proteins and disruption of cellular membranes, leading to localized tissue damage and triggering further biochemical responses aimed at repair or defence*.

The secondary phase of impedance changes across the entire leaf signifies a systemic response that integrates local damage signals into broader physiological adaptations. This dynamic bears resemblance to stress responses in non-excitable animal cells, where localized injuries also trigger cascades of signaling events leading to alterations in membrane potential and cellular metabolism^55^

#### Common features and differences from the above cases

In both plant and animal cells, these responses exemplify how living systems utilize electrical properties and biochemical signaling to adapt to environmental stresses while maintaining overall structural integrity.For instance, during ischemia, tissues experience an acute phase where metabolic processes are disrupted, followed by a delayed phase characterized by inflammation and tissue remodeling. This similarity suggests that organisms—whether plants or animals—exhibit common adaptive mechanisms when faced with external stresses. However, while there are overarching principles governing these responses, each type of stress may elicit unique physiological pathways due to differences in tissue structure and function.

Tumors often exhibit heterogeneous responses to therapeutic interventions; understanding how they react to stress can inform treatment strategies, e.g. if tumor exhibits a strong initial antiphase reaction to therapy, it may indicate an immediate defense mechanism that could lead to treatment resistance. Amplitude of the secondary reaction could provide insights into how well tumor adapts over time to therapeutic pressures, potentially guiding decisions on whether to continue or modify treatment protocols. Thus, assessment of these biphasic responses is critically important when considering methods for oncology treatment and tumor reprogrammin^42,56.^

